# Resolving transcriptional states and predicting lineages in the annelid *Capitella teleta* using single-cell RNAseq

**DOI:** 10.1101/2020.10.16.342709

**Authors:** Abhinav Sur, Néva P. Meyer

## Abstract

Evolution and diversification of cell types has contributed to animal evolution. However, gene regulatory mechanisms underlying cell fate acquisition during development remains largely uncharacterized in spiralians. Here we use a whole-organism, single-cell transcriptomic approach to map larval cell types in the annelid *Capitella teleta* at 24- and 48-hours post gastrulation (stages 4 and 5). We identified eight unique cell clusters (undifferentiated precursors, ectoderm, muscle, ciliary-band, gut, neurons, neurosecretory cells and protonephridia), thus helping to identify previously uncharacterized molecular signatures such as novel neurosecretory cell markers. Analysis of coregulatory programs in individual clusters revealed gene interactions that can be used for comparisons of cell types across taxa. We examined the neural and neurosecretory clusters more deeply and characterized a differentiation trajectory starting from dividing precursors to neurons using Monocle3 and velocyto. Pseudotime analysis along this trajectory identified temporally-distinct cell states undergoing progressive gene expression changes over time. Our data revealed two potentially distinct neural differentiation trajectories including an early trajectory for brain neurosecretory cells. This work provides a valuable resource for future functional investigations to better understanding neurogenesis and the transitions from neural precursors to neurons in an annelid.

## Introduction

Proper development of multicellular organisms relies on precise regulation of the cell cycle relative to establishment of cell lineages and cell fate decisions, e.g., the maintenance of proliferating cells versus the onset of differentiation. In general, many embryonic and postembryonic tissues are generated by stem cells that give rise to multipotent precursor cells whose daughters differentiate into tissue-specific, specialized cell types. Cell fate acquisition and differentiation are directly regulated by changes in transcriptional gene regulation. Therefore, understanding the underlying transcriptional dynamics is of utmost importance to understand developmental processes. Furthermore, alterations in gene regulatory networks (GRNs) may have driven diversification of cell types during animal evolution. According to Arendt (2016), cell types are evolutionary units that can undergo evolutionary change. Therefore, to identify related cell types across taxa, it is necessary to compare genomic information, such as shared gene expression profiles or shared enhancers across individual cells from specific developmental regions and stages.

More recently, single-cell RNA sequencing (scRNAseq) has emerged as a powerful technique to understand the genome-wide transcriptomic landscapes of different cell types (Tang et al., 2010; Hashimshony et al., 2012; Saliba et al., 2014; Trapnell et al., 2014b; Achim et al., 2015; Satija et al., 2015; Kaia Achim, 2017; Vergara et al., 2017; Svensson et al., 2018; Zhong et al., 2018). scRNAseq enables massively parallel sequencing of transcriptomic libraries prepared from thousands of individual cells and allows for *in silico* identification and characterization of distinct cell populations (Trapnell, 2015; Tanay and Regev, 2017). It can therefore provide information regarding the various cell types that emerge during developmental processes (e.g. neurogenesis) and elucidate how the transcriptomic landscape changes within stem cells and their progeny as development progresses. As scRNAseq analysis algorithms allow for *a priori* identification of individual cells within a population, one can process heterogenous cell populations and unravel the transcriptomic signatures underlying such heterogeneity. This allows for discovery of novel cell types and resolution of the transcriptional changes throughout a single cell type’s developmental journey. Emergence of this technology has therefore made it possible to predict molecular trajectories that underlie cell fate specification by sampling across a large number of cells during development and connecting transcriptomes of cells that have similar gene expression profiles (Farrell et al., 2018). Such approaches have recently gained prominence in evolutionary developmental biology and are being used to understand evolutionary relationships between cell types across taxa. This has paved the way to systemic molecular characterization of cell types and developmental regulatory mechanisms in understudied metazoan lineages.

Although whole-organism scRNAseq approaches have been used to unravel cell type repertoires in animal clades outside and across Bilateria (Kaia Achim, 2017; Farrell et al., 2018; Plass et al., 2018; Sebe-Pedros et al., 2018a; Sebe-Pedros et al., 2018b; Foster et al., 2020), there has been limited systemic information regarding cell type diversity and regulatory mechanisms underlying differentiation trajectories in the third major clade Spiralia (≈Lophotrochozoa)(Marletaz et al., 2019). Whole-body scRNAseq has been performed on a few spiralians such as the planarian *Schmidtea mediterranea* (Cao et al., 2017; Plass et al., 2018) and the annelid *Platynereis dumerilii* (Achim et al., 2015; Kaia Achim, 2017). In *S. mediterranea*, different classes of neoblasts and various differentiation trajectories emanating from a central neoblast population were detected using whole-body scRNAseq (Plass et al. 2018). In *P. dumerilii* larvae, whole-body scRNAseq yielded five differentiated states — anterior neural domain, gut, ciliary-bands, an unknown cell population and muscles (Kaia Achim, 2017). Similar transcriptomic information from other spiralian taxa can provide insight into conserved cell types and their evolution.

In this manuscript, we used scRNAseq to characterize larval cell types at 24- and 48-hours post gastrulation in the annelid *Capitella teleta* (Blake A. J, 2009)(Fig. 1), highlighting potential genetic regulatory modules and differentiation trajectories underlying different cell types. We (i) classified the captured cells into several molecular domains, (ii) predicted lineage relationships between neural cells in an unbiased manner, and (iii) identified neurogenic gene regulatory modules comprising genes that are likely involved in programming neural lineages. We compared larval cell types identified in this study with those in *P. dumerilii* at roughly similar stages during development. This study provides a valuable resource of transcriptionally distinct cell types during *C. teleta* larval development and illuminates the use of scRNAseq approaches for understanding molecular mechanisms of larval development in other previously understudied invertebrates.

**Figure 1:**
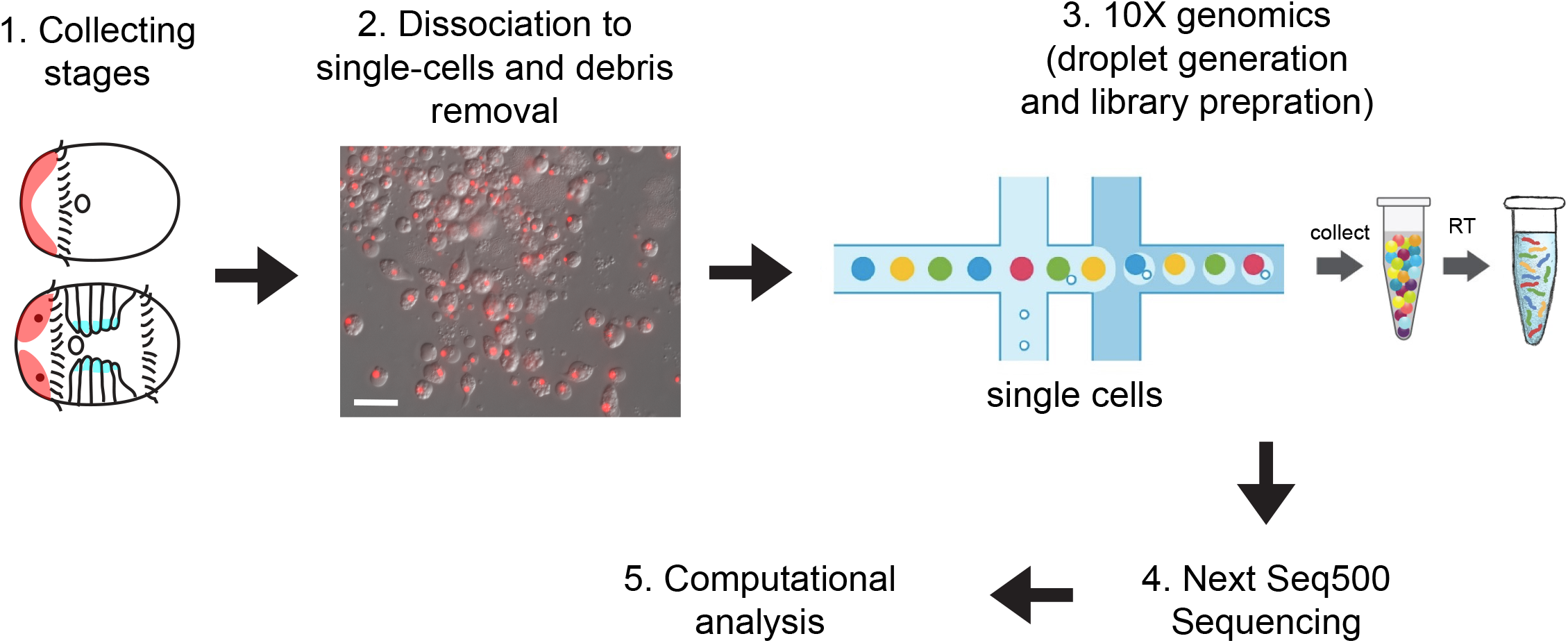
Single-cell transcriptomics of stage 4 and 5 *C. teleta* larvae. Whole-body stage 4 and 5 (1) larvae were dissociated into single cells using a combination of mechanical and enzymatic dissociation (2). Individual cells were randomly selected for droplet generation using the 10X Genomics Chromium Platform (3). Single-cell transcriptomes were pooled and sequenced using NextSeq500 High-output method (4) generating 573 million reads across the two samples. Sequences obtained were curated and aligned to the *C. teleta* genome v1.0 followed by application of downstream computational pipelines for clustering, trajectory analysis, pseudotime analysis and estimating RNA dynamics (5). Scale bar – 10 μm.

## Materials and Methods

For the data reported here, cell dissociation and scRNAseq using the 10X genomics platform was performed at the Single-cell Sequencing Core at Boston University, Boston, Massachusetts. We tried replicating the 10X experiment at the Bauer Sequencing Core, Harvard University; however, that run did not yield enough RNA for amplification and sequencing.

### *Capitella teleta* cell dissociation and single-cell suspension

Total number of cells in *C. teleta* larvae at stages 4 and 5 were estimated by counting Hoescht-labeled nuclei in the episphere at stages 4–5 (Fig. S1A, B) and from previously collected cells counts in the trunk at stages 4 and 5 (Sur et al., 2020). At both stages, the episphere was divided into 10 μm thick z-stacks and Hoecht^+^ nuclei were counted in each z-stack using the Cell Counter plugin in Fiji. At stage 4, Hoescht^+^ nuclei in the unsegmented trunk were counted within the presumptive neuroectoderm using a strategy previously described. Once the trunk neuroectoderm becomes segmented by stage 5, Hoescht^+^ nuclei were counted in segments 2–4 and 5–7 within specific region of interests (Sur et al., 2020). We also optimized cell-dissociation protocols in *C. teleta* (final protocol detailed below). Based on cell-dissociation trials using three different proteolytic enzymes (papain, trypsin and pronase), we found 1% papain yielded the highest number of dissociated cells but also led to a lower proportion of viable cells (Fig. S1C). Cell counts were estimated after mechanical and proteolytic digestion, size-exclusion of >40 μm cells or cell-clumps, and three washes in artificial seawater or cell-media (see below). Papain does not readily dissolve in seawater and hence needs to be resuspended in dimethylformamide or dimethyl sulfoxide, which may have an adverse effect on the viability of the dissociated cells. Cell dissociation using 1% Trypsin yielded the greatest number of viable cells and was used to dissociate *C. teleta* cells for scRNAseq (Fig. S1C).

Next, for collecting high-quality starting material for scRNAseq, healthy males and females were mated under controlled conditions and their offspring collected at the gastrula stage (stage 3) (Seaver et al., 2005; Sur et al., 2017). Stage 4 and stage 5 larvae were collected from two different sets of parents, each from a single mother. *Capitella teleta* embryos and larvae were incubated in artificial seawater (ASW) with 50 ug/mL penicillin and 60 ug/mL streptomycin at 19 °C for 1–2 days until they reached stage 4 prototroch or stage 5 (Fig. 1). For single-cell dissociation, 300 larvae from a single brood (i.e. from one male and female) for each stage were then collected into 1.5 mL centrifuge tubes and equilibrated in Ca^2+^/Mg^2+^-free ASW (CMFSW). Most of the CMFSW was removed from the tubes, and larvae were mechanically homogenized using separate, clean and sterile pestles as well as a hand-held homogenizer (Cole Parmer, LabGEN 7B) for 5–10 seconds. Homogenized larvae from each stage were then incubated in 1% Trypsin (SigmaAldrich, Cat# T4799-5G) in CMFSW for 30 minutes at room temperature with constant rocking. During incubation, dissociated tissues were periodically triturated using both wide-mouthed and narrow-mouthed Pasteur pipettes. After 30 minutes of incubation, the tissue lysate was passed through a 40 μm nylon cell-strainer (Fisherbrand, Cat# 22-363-547) to get rid of undissociated cell-clumps. The resultant cell-suspension was then centrifuged at 1100 x g for 7 minutes with slow-braking and washed twice in cell media that was developed originally for marine hemichordate cell-cultures (3.3X Dulbecco’s PBS and 20 mM HEPES, pH = 7.4; Paul Bump, Lowe lab, personal communication). Dissociated cells were then resuspended in the cell media and checked under an inverted phase-contrast microscope to ensure a single-cell suspension was obtained.

Cells were counted using a Neubauer hemocytometer, and survivability was assayed using a Trypan blue exclusion test. Cells were observed under the 20X objective of a Zeiss AxioObserver-5 inverted microscope following Trypan-blue staining available at the Boston University Single-Cell Sequencing Core to ensure cell viability prior to droplet generation. Previous practice dissociations of *C. teleta* larvae and visual inspection using a Zeiss M2 microscope at 40X resolution revealed dissociated cells ranging from 2–12 μm in diameter (Fig. 1). Due to the small size of *C. teleta* cells and the unavailability of a high-resolution microscope, the total number of cells dissociated in the cell-suspension could not be quantified confidently at

Boston University. Therefore, based on cell counts using the Zeiss AxioObserver-5 inverted microscope under the 20X objective, cells were diluted to a target of 400 cells/μL. However, due to our inability to accurately quantify cells and based on results from previous dissociation trials and total number of cells estimated per stage 4 and 5 larvae, our final cell suspension may have contained in the range of ~4000 cells/μL for stage 4 and ~2000 cells/μL for stage 5. A total of 15 μL of the resuspended cell-suspension was used for droplet generation estimating a 67% efficiency in droplet capture as per 10X genomics standard guidelines.

### Cell capture and sequencing

*C. teleta* larval cells were captured in droplets and run on the 10X genomics scRNAseq platform at the Boston University Single Cell Sequencing Core following the manufacturer’s instructions (Single Cell 3’ v3 kit). The cDNA library and final library after index preparation were checked with bioanalyzer (High Sensitivity DNA reagents, Agilent Technology #5067-4626; Agilent 2100 Bioanalyzer) for quality control (Fig. S1D). Following library preparation, sequencing was performed with paired-end sequencing of 150 bp each end on four lanes of NextSeq500 per sample using the Illumina NextSeq500 High-Output v2 kit generating ~573 million reads in total.

To rectify our inability to control the number of cells input in the first trial, we repeated the cell dissociation and cell-capture procedures at the Bauer Sequencing Core, Harvard University. In this trial, we carefully counted and diluted cells to 400 cells/μL under a Zeiss M2 microscope using a 40X objective and loaded 15 μL of the resuspended cell-suspension aiming to capture ~4000 cells per stage estimating a 67% efficiency in droplet capture as per 10X genomics standard guidelines. However, in this second trial, ~4000 captured cells did not generate sufficient cDNA yield following whole transcriptome amplification to enable library preparation for sequencing (Fig. S1E). This indicates that our first 10X genomics trial at Boston University probably represents the best quality output possible using the 10X genomics scRNAseq platform on *Capitella teleta* larval cells.

### Bioinformatic processing of raw sequencing data

Transcriptome sequencing analysis and read mapping were performed using CellRanger 2.1.0 according to the manufacturer’s guidelines. Reads were mapped onto the *Capitella teleta* genome v1.0 obtained from gene-models deposited at GenBank (GCA_000328365.1_Capca1_genomic.fasta; https://www.ncbi.nlm.nih.gov/assembly/GCA_000328365.1/) and Ensembl (Capitella_teleta.Capitella_teleta_v1.0.dna_sm.toplevel.fa; ftp://ftp.ensemblgenomes.org/pub/metazoa/release-46/fasta/capitella_teleta) using standard CellRanger parameters. The gene annotation files (.gff) files were downloaded from the respective genome databases. However, as CellRanger cannot read.gff files, each .gff file was converted into .gtf files using the gffread command from the cufflinks package (http://cole-trapnell-lab.github.io/cufflinks/file_formats/). Read mapping to the *Capitella teleta* genome v1.0 was visualized using the IGV 2.8.0 viewer. Mapping of the sequence reads to both Ensembl and GenBank sequences yielded similar results. CellRanger generated a Digital Gene Expression (DGE) matrix with genes as rows and cells as columns where paired-end reads, one containing the cellular and molecular barcodes (Unique molecular identifiers, UMIs) and the other containing the captured RNA fragment, were joined together in a .bam file and sorted using samtools. Reads already tagged with the cell and molecular barcodes (UMIs) were further trimmed at the 5’ end to remove Illumina-specific sequencing adapter sequences and at the 3’ end to remove poly-A tails using CellRanger default parameters.

### Gene annotation

To annotate the genes from the two versions of the *C. teleta* genome (Simakov et al., 2013), reciprocal BLAST comparison of individual gene sequences against the Swiss-Prot database was performed. For each transcript, the BLAST hit with the highest E-value was selected for annotation. The translated reference transcriptome along with the *C. teleta* gene-models were scanned using the HMMER suite 3.3 program hmmscan using default settings. Using HMMER and Pfam v31.0 database, protein domains in the *C. teleta* transcriptome were identified (Finn et al., 2016).

### Gene and cell filtering: Quality control and clustering analysis

DGE matrices were analyzed using the R package Seurat 3.1.4 (Satija et al., 2015).

Because of our inability to control the number of cells that were used for droplet generation, and to understand cell-state specific gene and UMI metrics, we performed an initial cluster analysis using less stringent gene and UMI cutoffs. Initially, gene per cell cutoffs between >200 and <3000 and UMI per cell cutoffs of >200 and <4000 were set. Genes that were expressed in at least three cells were kept and cells that had more than 5% mitochondrial reads were excluded. High mitochondrial content may indicate that a cell was stressed or dying. However, using such cutoffs, not enough unique cell clusters were detected. This may be because particular cell-doublet categories were not excluded in the cut-off selection. Therefore, this preliminary analysis led us to increase the gene per cell cutoffs to >300 and <2500 in order to prevent the inclusion of cell-doublets in our analysis.

We refined our gene/UMI cutoffs, and only genes that were expressed in at least three cells with a minimum of 300 genes were included in the analysis. Moreover, we also discarded cells with more than 2500 genes in sequences obtained from both samples in order to screen out cell-doublets. A total of 9487 genes across 1072 cells for stage 4 and 13403 genes across 1785 cells for stage 5 from one 10X genomics experiment were included in the final analysis. This accounts for around ~4 cells per stage 4 larva dissociated and ~6 cells per stage 5 larva dissociated that were bioinformatically recovered from initially loaded ~200 cells per larva for stage 4 and ~100 cells per larva loaded for stage 5. We only used 3.1% of Cell Ranger predicted captured cells for stage 4 and 10.86% of predicted captured cells at stage 5 for downstream analysis. UMI counts per gene in individual cells were normalized to the total UMI count of each cell using the ‘LogNormalize’ function with a scale factor of 10,000 (Fig. S2). Clustering analysis of the cells was done using the top 2000 variable genes identified using the ‘FindVariableFeatures’ function (selection.method = vst, nfeatures = 2000) (Fig. S2A, B). Following variable gene selection, data were then centered and scaled using the ‘Scale Data’ function with default parameters. These variable genes were then used to perform a principal component analysis (PCA) on the scaled data. The top 15 PCs obtained were then tested for significance using a JackStraw test that is part of the Seurat 3.1.4 parlance with 100 replicates. Principal components (PCs) with a p-value of less than 1e-5 were used to perform a Louvain-based clustering on the shared nearest neighbor (SNN) graph (Fig. S2E, F). For data visualization, we performed t-distributed stochastic neighbor embedding (t-SNE) and Uniform Manifold Approximation and Projection (UMAP) analysis. Specific cell-clusters were detected using the ‘FindClusters’ function from Seurat using a resolution of 0.5. Dendrograms depicting relationships between cell-clusters were generated using the ‘PlotClusterTree’ function.

Marker genes for individual clusters were identified using Seurat’s ‘FindAllMarkers’ function calculated using the Wilcoxon’s rank sum test. Using this approach, cells from each population were compared against each of the other clusters in order to detect uniquely expressed genes. Only genes that were enriched and expressed in 25% of the cells in each population (min.pct = 2.5) and with a log fold difference larger than 0.25 (logfc.threshold = 0.25) were considered. These differentially expressed genes per cluster were plotted on the feature plot individually using the ‘FeaturePlot’ function in Seurat for visualization in either UMAP or t-SNE space. The results also were visualized in a heatmap generated using the ‘DoHeatMap’ function.

### SWNE analysis

Apart from t-SNE and UMAP analysis, we also performed similarly weighted non-negative embedding (SWNE) analysis for visualizing high-dimensional single-cell gene expression datasets for each of our samples. SWNE captures both local and global structure in the data unlike t-SNE and UMAP embeddings, while enabling genes and biological factors that separate cell types to be embedded directly onto the visualization. To perform the SWNE analysis, a previously published R-based SWNE framework was used (Wu et al., 2018). The analysis was performed on log-normalized read count data for a set of variable genes from a previously generated Seurat object using the ‘RunSWNE’ function. Because the number of factor embeddings representative of the dataset cannot be estimated *a priori*, this parameter (called K) needs to be determined empirically. As the SWNE algorithm has the non-negative matrix factorization (NMF) inherently built into it, we initially performed NMF analysis over a broad range of K values ranging from 2 to 60 with steps of 2. The outputs of these separate runs were compiled together in order to estimate the optimal K-value. To find the optimal number of factors to use, the ‘FindNumFactors’ function was used. The function iterates over multiple values of k and provides the optimal number of factors that best represent the dataset. Following that, the NMF decomposition was run using the ‘RunNMF’ function that generates an output of gene loadings (W) and NMF embeddings (H). Following the NMF analysis, the SWNE embedding was run using the parameters: alpha.exp = 1.25, snn.exp = 0.25 and n_pull = 3 that control how the factors and neighboring cells affect the cell coordinates. The SWNE output was analyzed using the gene loadings matrix. Since NMF creates a part-based representation of the data, the factors often correspond to key biological processes or gene modules that explain the data. The top factors for each gene were visualized as a heat map using the ‘ggHeat’ function.

### Subclustering of neural cells

The neural and neurosecretory clusters obtained in the stage 5 t-SNE plot were isolated from the differential gene expression matrix, and the previously described Seurat analysis was repeated with the clustering resolution set at 0.5.

### Monocle3 pseudotime analysis

Pseudotime analysis of the neurogenic lineage was performed using the Bioconductor package Monocle3.0.2 (Trapnell et al., 2014b). For pseudotime analysis, the previously used Seurat object generated from the neural cell subcluster was imported into Monocle3. Monocle3 was run on our normalized counts matrix for the subclustered neural dataset. The data was subject to UMAP dimensional reduction and cell clustering using the ‘cluster_cells’ function (‘cluster_cells’: resolution=0.001). A principal graph was plotted through the UMAP coordinates using the ‘learn_graph’ function that represents the path through neurogenesis. This principal graph was further used to order cells in pseudotime using the ‘ordercells()’ function in Monocle3. Following that, we identified the population of neural precursor cells (NPCs) based on expression of cell-cycle markers and re-ran ‘ordercells()’ with NPCs as the root cell state. Genes changing as a function of pseudotime along the principal graph were determined using ‘graph_test’ function. Cells and most differentially expressed genes were then plotted in pseudotime using default parameters in Monocle3. The most significantly expressed genes with the greatest q-values were plotted on a heatmap of expression over pseudotime using the ‘plot_pseudotime_heatmap’ function in Monocle.

### RNA velocity estimation

To calculate RNA velocity of single cells within the neural subcluster, we applied the velocyto R (v0.6) package (La Manno et al., 2018). Velocyto uses the mapped reads from CellRanger and counts the number of spliced and unspliced reads separately. As the CellRanger read-mapping algorithm is splice-sensitive, the RNA velocity analysis can very easily be applied on the .bam files generated by CellRanger. For our 10X output, counting was performed at the level of molecules, taking into consideration the annotation (spliced, unspliced etc.) of all reads associated with the molecule. A molecule was annotated as spliced, unspliced or ambiguous based on the following criteria: a molecule was considered spliced if all of the reads in the set mapped only to the exonic regions of the compatible transcripts whereas a molecule was called as unspliced if at least one of the supporting reads were found to span exon-intron boundaries or mapped to the intron of the transcript. Molecules for which some of the reads mapped exclusively to the exons and some exclusively to the introns were categorized as “ambiguous” and not used for downstream analysis (La Manno et al., 2018). The command-line interface (CLI) for velocyto R (v0.6) was run in permissive mode. In this setting, we only used the cells mapped to the transcriptome that were present in our final neural subclustering Seurat analysis. Using all cells from the stage 5 neural subcluster, we normalized the expression per cell and selected the top 2000 variable genes to perform a PCA. Using the first 15 principal components we performed a data imputation with a neighborhood of 200 cells (k = 200 nearest neighbors) and calculated RNA velocities. All steps were performed using in-built parameters for fitting gene-models, predicting velocity, extrapolating and plotting. To visualize the plots, we used the t-SNE embedding as produced by the Seurat analysis.

## Results

### Single-cell profiling of *C. teleta* stage 4 and 5 whole-body larvae

To explore developmental trajectories and how transcriptomic landscapes across cells change during early larval development in the marine annelid *Capitella teleta*, we dissociated 300 whole larvae from a single brood at both 24- and 48-hours post gastrulation, which corresponds to stage 4 just after appearance of the prototroch ciliary band, and stage 5, respectively. Based on Hoescht-labeled nuclei counts in the episphere and the trunk, we estimated that a stage 4 larva has ~2000 cells and a stage 5 larva has ~4000 cells (Fig. S1A, B). To enable random sampling of cells, we used 300 animals per stage to maximize the initial pool of cells for cell-capture. We also tested multiple methods of cell dissociation and examined cell survival rate (Fig. S1C). For scRNAseq, we dissociated cells in 1% trypsin for 30 minutes since this yielded the best survival rate (97%). However, due to the unavailability of a high-resolution microscope at the genomics facility and the small size of *C. teleta* dissociated cells (2–12 μm), total number of cells could not be counted accurately (See Materials and Methods). Therefore, based on previous pilot cell-dissociation trials (Fig. S1C), the number of cells were roughly estimated in the dissociated cell-suspension. We intended to sequence ~4000 cells per stage but due to technical limitations, we estimate that a much higher concentration of cells (see Methods) was loaded into the droplet-based scRNAseq platform 10X Genomics Chromium (Fig. 1). After sequencing and read-mapping, CellRanger predicted to have recovered 34,592 cells with 7,251 mean reads/cells from stage 4 and 16,434 cells with 17,837 mean reads/cell from stage 5 (Table S1). Based on our rough estimations, we recovered around 55% of the total number of cells input into the 10X Genomics system (~60,000 for stage 4 and ~30,000 for stage 5). However, there appeared to be a lot of noise due to the presence of cell-doublets and free-flowing RNA following cell-capture and sequencing. Hence, to identify distinct cell types from the stage 4 and 5 single-cell datasets, the assembled reads were passed through stringent Seurat quality control and UMI filtering algorithms (Fig. S2). A second trial conducted with careful estimation of cell counts and capturing ~4000 cells/stage did not yield enough cDNA to make high quality sequencing libraries unlike the first trial (Fig. S1D, E). Following computational filtering of our dataset to remove low-complexity transcriptomes, lowly-expressed genes and transcriptome doublets, we bioinformatically recovered 1072 cells from 300 stage 4 larvae and 1785 cells from 300 stage 5 larvae that were used for downstream analysis. Although we only captured a small fraction of cells after computational filtering, this is the first ever scRNAseq experiment on *C. teleta* larvae using the 10X genomics platform, and we were able to resolve some discrete transcriptional profiles and their underlying developmental trajectories.

### Transcriptional cell states in stage 4 and 5 larvae

To classify cell population identities in the global dataset across the two *C. teleta* larval stages, Seurat unsupervised clustering (Butler et al., 2018) of the aggregated data from stages 4 and 5 was conducted. UMAP analysis revealed six computationally identified clusters with a tight group of cells (C0, C1, C3, C4, and C5) and one cell-population situated farther away (C2; Fig. 2A). As at stages 4 and 5, majority of the cells in the *C. teleta* body are undifferentiated, and these individual clusters likely represent distinct developmental trajectories through which cells are progressing. Undifferentiated cells expressing receptors of growth factors (e.g. *fgfrl1, egf-like receptors*) and cell-cycle regulatory genes (e.g. *cdc6, mcmbp, cks1*; Fig. 2B, C) were found to be scattered across all cell clusters. In order to assign cluster identity, we used previously characterized genes in *C. teleta* and uncharacterized *C. teleta* genes homologous to known tissue markers in other taxa. Each cluster was identified based on the analysis of the top 30 significantly enriched genes per cluster. Gene annotations are reported in Table S2.

**Figure 2:**
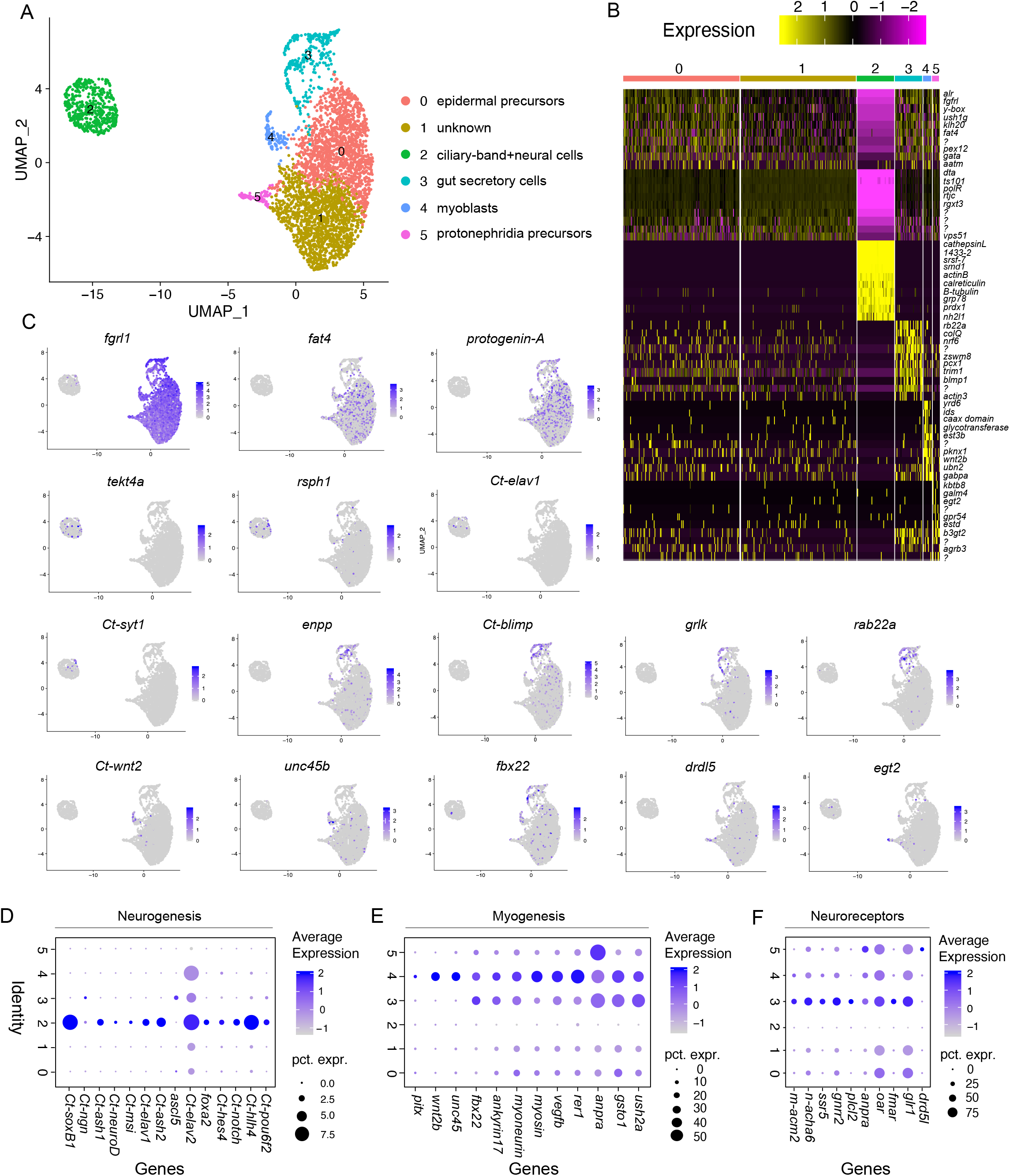
Mapping of *C. teleta* larval tissues from aggregated stage 4 and 5 datasets. (A) UMAP representation of the aggregated data (stages 4 and 5), where the clustering of cells depicts their transcriptional similarity. (B) Heatmap of the top 10 genes significantly enriched in each cluster. Representative gene names obtained from closest reciprocal BLAST hits are shown close to each row. The full gene-list is in Tables S2. (C) UMAP plots showing log-normalized counts of select representative genes from each cluster. Color intensity is proportional to the expression level (purple: high; grey: low). (D) Dotplot of representative genes involved in *C. teleta* neurogenesis. (E) Dotplot showing novel markers implicated in *C. teleta* myogenesis. (F) Dotplot showing orthologs of neurotransmitter and neuropeptide/neurohormone receptors across clusters.

Cluster C0 was enriched in genes predicted to be involved in extracellular matrix remodeling such as *protogenin-A, protocadherin fat-4*, tyrosine protein kinase *csk1, chaoptin* and *hepatocyte growth factor (hgf)*, indicating that these cells may be epidermal precursor cells (Fig. 2B, C). Similarly, differentially expressed genes in the C1 cluster included *UDP-D-xylose:L-fucose alpha-1,3-D-xylosyltransferase 3 (rgxt3), D-threonine aldolase (dta)*, a chitin-binding peritrophin-A domain containing protein, and vacuolar protein sorting-associated protein 51 homolog (*vps51*) (Fig. 2B, C), all of which represent chitin-binding proteins and proteoglycans (Shen and Jacobs-Lorena, 1999); however, the exact identity of cells in this cluster remained unclear. The other clusters also had distinct expression profiles suggestive of specific identities, C2: ciliary bands + neural cells (*tekt4a, rsph1, Ct-elav1, Ct-syt1*) among others, C3: gut secretory cells (*colq, Ct-blimp, glna2, enteric neuropeptides*), C4: myoblasts (*Ct-wnt2*, tetratricopeptide domain containing *unc45b*, F-box protein homolog *fbx22, vegfb, rer1, myosin heavy-chain*), and C5: protonephridia (*S-formylglutathione hydrolase, hercynylcysteine sulfoxide lyase and carbohydrate sulfotransferase 1*) (Fig. 2B–E). C3 also expressed some myogenic markers, albeit at a lower level (Fig. 2E), which could indicate that a subset of developing muscle precursors clustered here.

Interestingly, apart from C2, all other clusters expressed receptors for neurotransmitters and neurohormones (Fig. 2F). C5 (protonephridia) was found to express dopaminergic neuroreceptors (*drd5l*) and atrial-natriuretic peptide receptors (*anpra*) (Fig. 2F), while C3 was particularly enriched in receptors for neurotransmitters and neuropeptides/hormones like acetylcholine (*acm2* and *acha6*), GABA (*plcl2*), FMRF-amide (*fmar*), gonadotropin (*gnrr2*) and somatostatin (*ssr5*). Even though the C3 cluster contained cells that expressed neurotransmitter and neurohormone receptors, we think this cluster could largely contain gut and muscle cells based on expression of these types of receptors in these cell types in other taxa (Florey and Rathmayer, 1978; Walker et al., 1993; Terra et al., 2006; Crisp et al., 2010; Mirabeau and Joly, 2013; Hung et al., 2020; Wu et al., 2020a). Receptors for acetylcholine, FMRF-amide and GABA have been reported to be localized to the body wall muscle in earthworms and leeches (Walker et al., 1993). Spiralian FMRF-amide G-protein coupled receptors (GPCRs) were first reported in P. *dumerilii* and were found to be homologous to insect neuropeptide receptors responsive to neuropeptide-F (Elphick et al., 2018). In *C. teleta*, FMRF-amide^+^ neurons have been shown to be associated with the midgut (Meyer et al., 2015). Both glutamate and GABA signaling have been reported in midgut epithelial cells in insects (Terra et al., 2006; Hung et al., 2020). Somatostatin/allatostatin-C encodes for a neuropeptide family of hormones that are expressed in *D. melanogaster* midgut endocrine cells (Wu et al., 2020a), while octopamine GPCRs have been reported in the annelid *P. dumerilii* and the priapulid *Priapulus caudatus* where they were shown to be activated in presence of dopamine, tyramine and octopamine ligands (Bauknecht and Jekely, 2017). Such neuropeptide- and neurotransmitter-signaling repertoires may regulate diverse behavioral changes associated with life-phase transitions in *C. teleta* based on previous evidence from *P. dumerilli* (Conzelmann et al., 2013).

The neurotransmitter and neuropeptide receptors characterized in our dataset can also serve as a valuable resource for better understanding neurotransmitter and neuropeptide signaling in *C. teleta*.

### Overall molecular changes across *C. teleta* larval development

An unsupervised graph-based clustering approach was used to separately analyze transcriptomic data at stages 4 and 5. Datasets were visualized with t-SNE dimensionality reduction (Fig. 2, Fig. 3A, Fig. 4A). In our stage 4 dataset, we detected ~174 median genes per cell and around ~740 median UMIs per cell, while in our stage 5 dataset, we detected ~241 median genes per cell and ~1145 median UMIs per cell (Fig. S2, Table S1). At both stages 4 and 5, t-SNE analysis revealed a large population of cells (C0; gray) that were enriched in ribosomal genes (RL10, RS9, RS4), cell proliferation markers (e.g. *pcna*), S-phase and M-phase cell-cycle markers (e.g. *cks1, mcm3, rfa3, wee1*) and chromatin remodeling genes (e.g. *acinu, bptf*), indicating that these cells are undifferentiated, developmental precursors (Fig. 3A, B, D, G, Fig. 4A, B, Table S3, S4). Such a finding is expected as the *C. teleta* larval body at this stage largely comprises proliferative cells (Seaver et al., 2005; Sur et al., 2020). The C0 cluster may contain different subsets of precursors or stem cells that give rise to different tissues throughout *C. teleta* development. Expression of *Ct-soxB1* in many of these cells indicates that at least a subset is ectodermal in origin (Fig. 3B, D, Fig. 4B, C). At stage 5 but not stage 4, some cells within C0 were also found to express muscle-associated markers such as *hand2, troponinC* and *twitchin* (data not shown). Therefore, we generically named this cluster ‘precursors’. In both datasets, we detected a few C0 cells that expressed *Ct-piwi1* and *Ct-hes2* (Fig. S3A, Fig. S5A). *Ct-piwi1* has been characterized as a marker of both somatic and germline stem-cells in *C. teleta* (Giani et al., 2011), while *Ct-hes2* is a homolog of the vertebrate *hes1a* gene, which is a *Notch* target and regulates stem-cell maintenance and cell-cycle progression. *Ct-hes2* is broadly expressed in larvae, including in lateral ectoderm and the posterior growth zone where new segments are generated from stage 7 onward in *C. teleta* (Thamm and Seaver, 2008). C0 in both datasets comprises a complex set of cells that express markers representative of various tissues and may represent uncommitted cells with different developmental trajectories.

**Figure 3:**
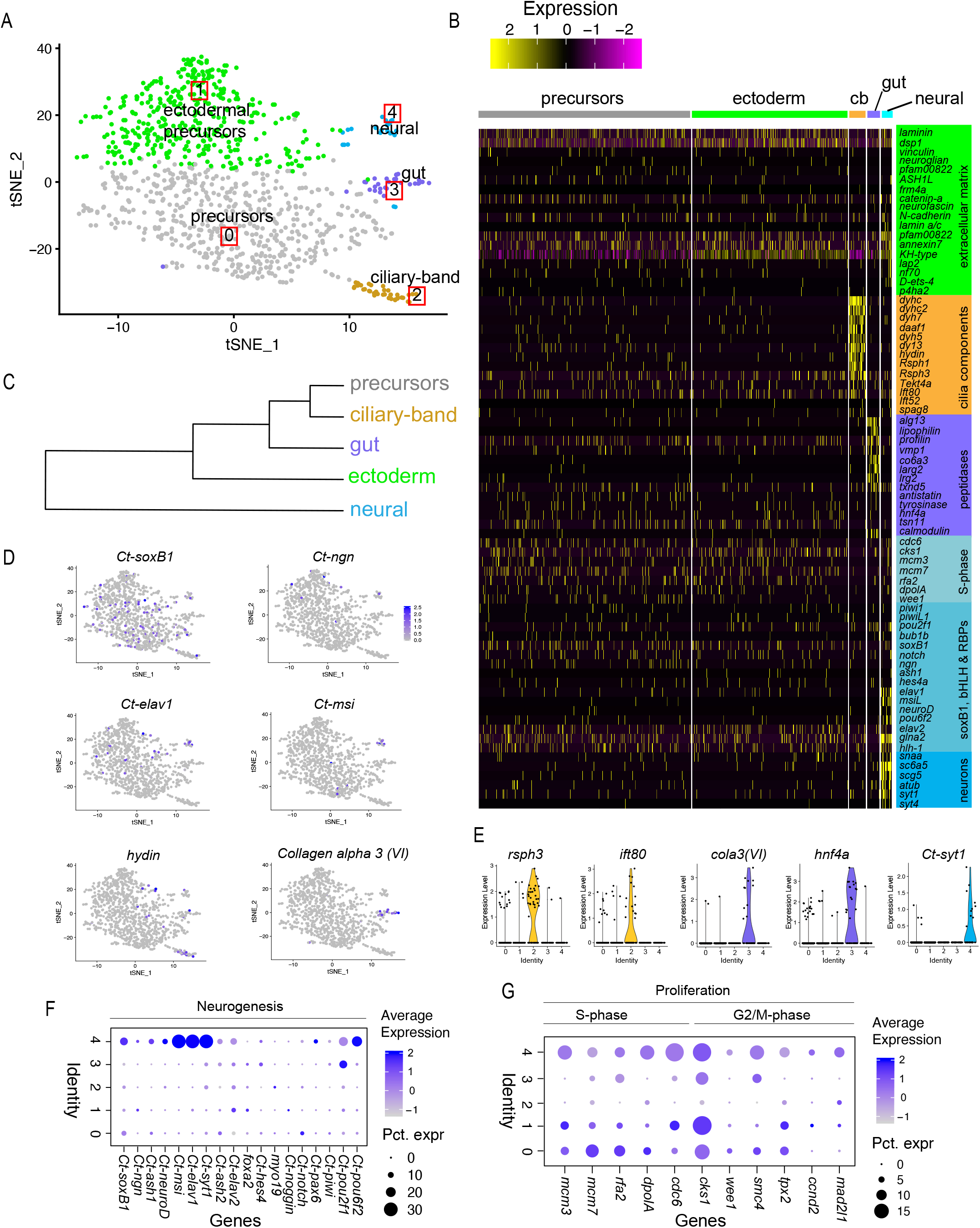
Single-cell molecular landscape of stage 4 *C. teleta* larvae. (A) t-SNE representation of stage 4 larval single cells with clusters labeled by molecular identities. (B) Cell type specific marker genes reflect cellular identities and functions. Heatmap showing log-normalized differentially expressed genes per molecular domain identified. Each row represents a single gene whereas each column represents a cell. (C) Analysis with PlotClusterTree in Seurat to reveal transcriptomic similarities between clusters. (D) t-SNE plots of cells colored by expression of selected marker genes that were used for identifying each molecular domain. The color key indicates expression levels (purple: high; grey: low). (E) Violin plots summarizing the expression levels of select representative genes per cluster. Data points depicted in each cluster represent single cells expressing each gene shown. (F) Dotplot of representative genes involved in *C. teleta* neurogenesis at stage 4. (G) Dotplot showing cell proliferation (S-phase and G2/M-phase) markers in the stage 4 clusters. cb: ciliary-band.

**Figure 4:**
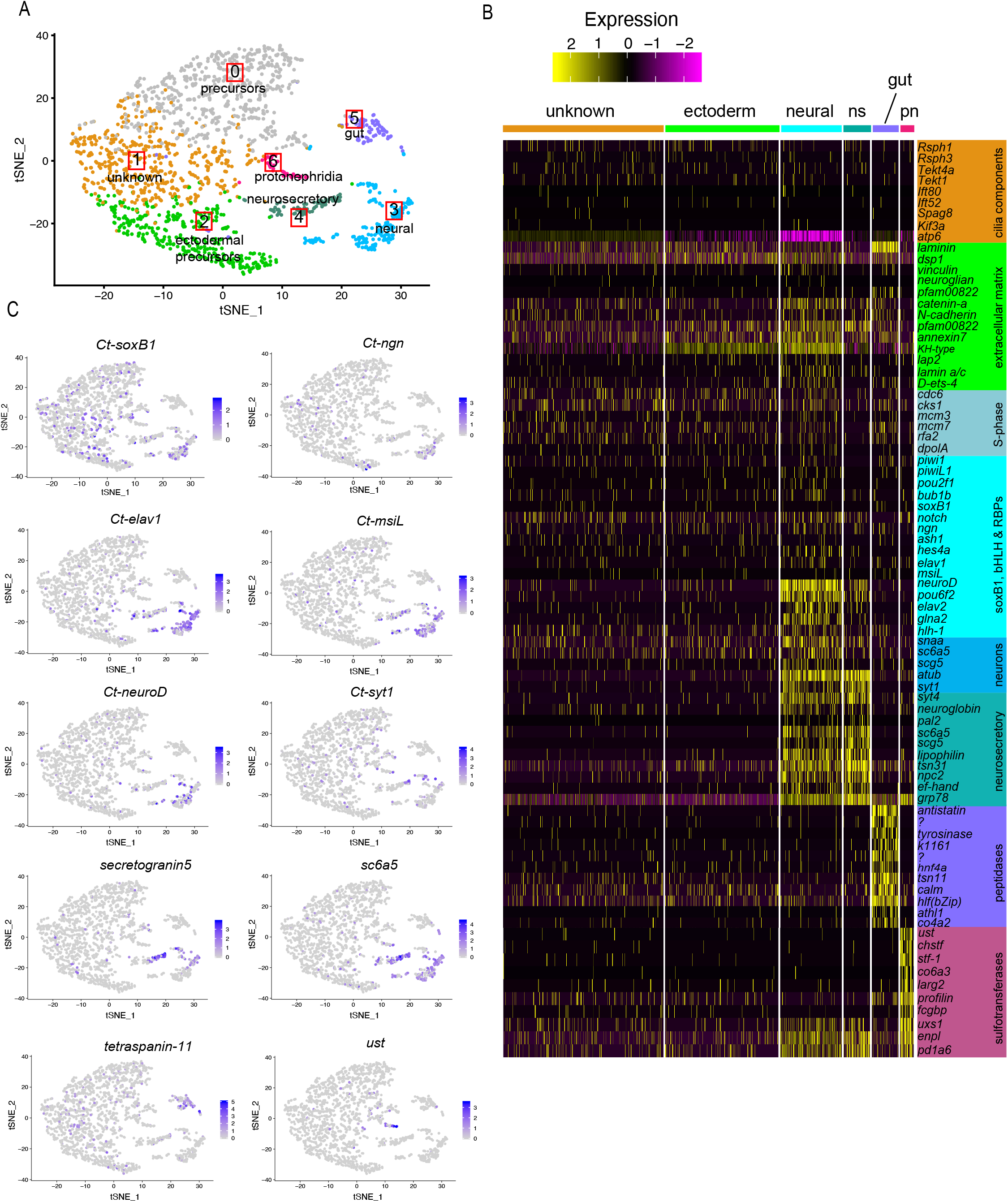
Single-cell molecular landscape of stage 5 *C. teleta* larvae. (A) t-SNE representation of stage 5 larval single cells with clusters labeled by transcriptional identities. (B) Cell type specific marker genes reflect cellular identities and functions. Heatmap showing log normalized differentially expressed genes per molecular domain identified. Each row represents a single gene whereas each column represents a cell. (C) t-SNE plots of cells colored by expression of selected marker genes that were used for identifying each cluster. The color key indicates expression levels (purple: high; grey: low). ns, neurosecretory; pn, protonephridia.

C1 in the stage 4 dataset and C2 in the stage 5 dataset primarily expressed markers associated with cellular tight junctions and extracellular matrix (e.g. *claudin, lamin a/c, annexin7* and *p4ha2*) as well as genes shared with C0 and the neural cluster (Fig. 3B, Fig. 4B, Fig. S3B, Fig. S5B). Lamin A/C and *p4ha2* have been identified in the epidermis of *P. dumerilii* and other spiralians (Kim et al., 2012; Kaia Achim, 2017). *Claudin* is a tetraspanning transmembrane protein that is an integral component of tight junctions (Krause et al., 2008; Piontek et al., 2008) while *annexin-7* has calcium-dependent membrane-binding activity in most animals. Finding shared expression of putative epidermal and neural markers could corroborate previous lineage tracing data suggesting that individual precursor cells in the neuroectoderm generate anywhere from one to 50 neural cells as well as one or two epidermal cells (Meyer and Seaver, 2009). Therefore, based on the expression of extracellular matrix remodeling genes and neural markers, C1 in stage 4 and C2 in stage 5 was identified as “ectodermal precursor cells”. Cells in the C1 cluster in the stage 5 dataset (Fig. 4A) expressed similar genes as in C1 of the aggregated dataset (Fig. 2A, B) such as *D-threonine aldolase* (*dta*) and vacuolar protein sorting-associated protein 51 homolog (*vps51*) as a result of which its identity remained unclear.

In the stage 4 dataset, C2 represents ciliary-band cells that express homologs in the *dynein* family (*dyhc, dyhc2, dyh7, dyh5, dy13, hydin* etc.), radial spoke-head genes (*rsph1* and *rsph3*) and intraflagellar transport-proteins (*ift80*) associated with the axonemal apparatus of cilia (Fig. 3B, E, Fig. S3C), similar to *P. dumerilii* (Kaia Achim 2017). Moreover, these ciliary-band cells at stage 4 do not express any of the S-phase or M-phase markers indicating that these cells are not proliferating (Fig. 3B, G). *Hydin* encodes for a protein that constitutes the axonemal central-pair apparatus that regulates cilia motility while IFT80 constitute part of the molecular machinery underlying cilia motility. However, at stage 5, cells expressing these same ciliary markers were found to be scattered across all clusters and did not resolve as a distinct cluster (Fig. 4B, Fig. S5C).

C3 in the stage 4 dataset (Fig. 3A, B) and C5 in the stage 5 dataset (Fig. 4A, B) were identified as “gut” based on some of the highly expressed markers in that cell-cluster including peptidases (e.g. *antistatin, tyrosinase*), secretory proteins (e.g. *lipophilin, profilin*) and glycotransferases (e.g. *lrg2b* and *alg13*) (Fig. 3B, Fig. 4B). These cells were also found to express *hepatocyte nuclear factor 4a (hnf4a), tetraspanin-11* and *collagen alpha* (Fig. 3B, D, E, Fig. 4B, C, Fig. S3D, Fig. S4D), which have been shown to be expressed in midgut cells in *P. dumerilii* (Kaia Achim 2017) and in digestive cells of the cnidarian *Nematostella vectensis*, the ctenophore *Mnemiopsis lyeidi* and the sponge *Amphimedon queenslandica* (Sebe-Pedros et al., 2018a; Sebe-Pedros et al., 2018b). However, *collagen alpha* expression was not detected in the stage 5 “gut” cluster while *tetraspanin-11* was one of the most enriched genes in the C5 cluster of the stage 5 dataset (Fig. 4C). Interestingly, *Ct-gataB1*, which is expressed in endodermal cells at stage 4 in *C. teleta* and in the large, yolky midgut cells at later larval stages, was found to be excluded from the ‘gut’ cluster at stage 4 but not at stage 5 (Fig. S3D). However, at both stages, *hnf4a* and *Ct-gataB1* were expressed in a subset of cells in the C0 “precursors” clusters (Fig. S3D, Fig. S5D). Previous lineage tracing experiments identified a population of small, interstitial cells in the midgut of *C. teleta* larvae (Meyer et al., 2010), but the genes expressed in these interstitial midgut cells have not been characterized. Since the C3 cluster at stage 4 expresses digestive enzymes but not *Ct-gataB1*, these could represent interstitial midgut cells. At stage 5, the C5 cluster may include both *Ct-gataB1^+^* large, yolky midgut cells as well as interstitial midgut cells. One possible reason for the clustering of *Ct-gataB1^+^* cells among precursor cells at stage 4 may be because of the proliferative nature of early endodermal cells. As a result, our bioinformatic pipeline detected these *Ct-gataB1^+^* cells to be more similar to the dividing precursors than the C3 gut cells. At stage 5, the *Ct-gataB1^+^* cells may have a decreased proliferative potential and hence are clustered with the other ‘gut’ cells. However, this awaits further verification using cell proliferation assays and in-situ hybridization.

Lastly, the C4 cluster in the stage 4 dataset and C3 in the stage 5 dataset likely have a neural identity based on expression of neural differentiation markers such as *Ct-elav1, Ct-syt1, Ct-msi, Ct-neuroD* and *Ct-syt1* (Meyer and Seaver, 2009; Meyer et al., 2015; Sur et al., 2017)(Fig. 3B, D–F, Fig. 4B, C, Fig. S3F, G, Fig. S5F, G). S-phase markers were also expressed in the ‘neural’ cluster at stages 4 and 5 indicating that these may be dividing neural progenitors given their spatial proximity to *Ct-elav1^+^* and *Ct-syt1^+^* cells in the tSNE plot (Fig. 3B, D, Fig. 4B, C, Fig. S3E, Fig. S5E). In the stage 4 dataset, a few cells in C1 were also found to express some neural markers (e.g. *Ct-ngn, Ct-ash1, Ct-msi* and *Ct-elav1*) (Fig. 3D). At stages 4 and 5, previous work using whole-mount in situ hybridization found that *Ct-ngn* and *Ct-ash1* are expressed in neural precursor cells (NPCs) and dividing foregut precursor cells. *Ct-ash1* is also expressed weakly in dividing mesodermal precursor cells and in some ectodermal cells outside the neuroectoderm (Meyer and Seaver, 2009; Sur et al., 2017; Sur et al., 2020). Therefore, in the C1 cluster at stage 4, the *Ct-ngn^+^/Ct-ash1^+^* cells may be ectodermal precursors, NPCs, and/or foregut precursor cells, while *Ct-ngn^-^/Ct-ash1*^+^ cells may be mesodermal precursor cells. Previous work has also shown that *Ct-msi*, *Ct-elav1* and *Ct-syt1* are exclusively expressed in differentiating and differentiated neurons (Meyer and Seaver, 2009; Meyer et al., 2015; Sur et al., 2017; Sur et al., 2020). Therefore, *Ct-elav1^+^/Ct-syt1^+^* cells in the C4 cluster at stage 4 are likely neurons, which first form in the developing brain and around the mouth (Meyer et al., 2015). At stage 5, the neural cells form a more coherent cluster C3, comprising NPCs expressing cell-cycle markers, intermediate differentiation states expressing *Ct-ngn, Ct-neuroD* and *Ct-ash1*, and mature neurons expressing *Ct-elav1, Ct-msi* and *Ct-syt1* (Fig. 4B, C; Fig. S5F, G). *Ct-elav1* and *Ct-syt1* were found to be more enriched and restricted to C3 in the stage 5 dataset unlike our observations at stage 4. At both stages we also observed the expression of *Ct-hunchback* in the neural cluster as previously reported in *Capitella* (Werbrock et al., 2001) and *P. dumerilii* (Kerner et al., 2006), and a homolog of MAP-kinase interacting serine/threonine kinase (*MKNK1*) in neural cells possibly indicating the involvement of the MAP-kinase signaling pathway during *C. teleta* neural development.

At stage 5, we also identified two additional discrete clusters from stage 4. C4 in the stage 5 dataset was classified as ‘neurosecretory’ based on the expression of markers genes such as the sodium- and chloride-dependent glycine transporter (*sc6a5*) and *synaptotagmin-4* (*syt4*) and *secretogranin-V*(*scg5*) (Fig. 4B, C, Fig. S5G, H). Neurosecretory cells secrete neuropeptides or hormones in response to neural input. A few neurosecretory cells were also detected as early as stage 4, and these cells clustered within the neural cluster (C4) (Fig. 3B; Fig. S3G, H). These cells could represent neurosecretory brain centers, which have been previously reported in other annelids (Tessmar-Raible et al., 2007; Williams et al., 2017). The function of the *sc6a5* gene is to impart neurosecretory fate by inhibiting glycinergic neurotransmission. In addition, non-calcium binding members of the Synaptotagmin family (i.e. Syt4) and *Syt-alpha* are implicated in the generation of large, dense-core vesicles for neurosecretion and have been found to be highly expressed in neurosecretory cells (Moghadam and Jackson, 2013; Park et al., 2014). *Secretogranin-V* is a neuroendocrine precursor protein that regulates pituitary hormone secretion in mammals. Marker genes that characterize the neurosecretory cell-cluster (C4) are largely expressed within the neural cluster (C3) as well in the stage 5 dataset. A few unique genes expressed by the neurosecretory cells were *neuroendocrine convertase 2, prohormone convertase* (Fig. S5H), and *conopressin/neurophysin* (data not shown). Neurophysin has been characterized in the developing neurosecretory brain centers in the annelid *P. dumerilii* and zebrafish *D. rerio* (Tessmar-Raible et al., 2007). In addition to neurosecretory cells, we also identified a protonephridia cluster, C6, at stage 5 based on the expression of sulfotransferases involved in the urea-cycle, e.g., *uronyl sulfotransferase, UDP glucouronic acid decarboxylase* and *carbohydrate sulfotransferases* (Fig. 4B, C).

We further examined relationships between all cell-clusters using PlotClusterTree in Seurat as this better represents transcriptional similarities between clusters than t-SNE distance. We found that the neural cluster branches out first followed by the other non-neural cell-clusters (Fig. 3C). This further confirms that the neural tissue is the first to be specified during *C. teleta* development and exhibits more transcriptional similarity to ectodermal precursor cells (C1) than any other cluster.

To decipher differentially-expressed, coregulatory gene modules within each cluster, we also projected both datasets using SWNE on a high-dimensional space correlated with non-negative matrix factorization (NMF) factor embeddings (Wu et al., 2018). In the stage 4 dataset, our SWNE visualization (Fig. S4) showed a central precursor population that branches into four differentiation trajectories: ectoderm, ciliary-band, foregut and neural (Fig. S4A, B). At stage 5, the protonephridia cluster emanated from the central ectodermal cluster while the neural cluster split into two, giving rise to the neurosecretory cluster (Fig. S6A, B).

Based on our SWNE embeddings plot, we deciphered differentially expressed genes for each cluster. The highest number of differentially expressed genes were found in the neural cell-cluster in both stages. For example, at stage 4, a glutamate receptor gene *grik4* and a tyrosine-protein phosphatase non-receptor type 4 gene (*ptn4*) were found to be coregulated together in a subset of neural cells (Fig. S4B). The gene *ptn4* encodes for a non-receptor tyrosine kinase (nRTK), and members of this family have been found to be abundantly present in excitatory synapses in the mammalian brain where they interact directly with glutamate receptors and phosphorylate tyrosine sites (Mao and Wang, 2016). Hence, cells expressing “factor 3” within the neural cluster may represent neurons that are excited by glutamate (Fig. S4A). These may also represent one of the first neuronal sub-types to differentiate during early *C. teleta* development. At stage 5, some uniquely expressed genes in the neurosecretory cells revealed by our SWNE analysis include *myom1* (myomodulin neuropeptides 1) and *orckB* (orcokinin neuropeptides class B) (Fig. S6B). Myomodulin is a bioactive neuropeptide that was found to be secreted by a cholinergic motor neuron in the mollusk *Aplysia californica* and regulates contraction of the buccal muscles during feeding (Cropper et al., 1987). Putative neurosecretory cells expressing *myom1* in *C. teleta* were coregulated with other G-protein coupled receptor messengers such as *plpr1* and *y1760* that provide insight into the *myom1* mediated neuropeptide signaling pathway (Fig. S6B). An orcokinin-like neuropeptide was previously identified in the *C. teleta* genome (Veenstra, 2011). Orcokinins have been detected in multiple other taxa such as insects, crustaceans, tardigrades, mollusks and sea-stars. In crustaceans, orcokinin neuropeptides have been shown to act as neuromodulators in the CNS and regulate peripheral neuromuscular junctions (Li et al., 2002). Using our unsupervised graph clustering and SWNE analysis, we show developmental trajectories of multiple cell types simultaneously, which was previously not possible using other techniques in *C. teleta*.

### Sub-clustering of neural cells reveals neural cell type diversity during neurogenesis

To gain better insight into the different neural cell types present in our stage 5 dataset, we further subclustered and curated cells from the neural (C3) and neurosecretory clusters (C4) using Seurat to obtain neural-specific t-SNE and UMAP plots (Fig. 5, Fig. 6, Fig. S7). Some proliferative cells expressing *Ct-soxB1* and bHLH factors like *Ct-ash1* and *Ct-ngn* were found to cluster within the C0 and C2 cells, however, their exact identity was not clear (see previous section) and hence these cells were not included in this analysis. As t-SNE plots do not preserve global data structure, i.e., only within cluster distances are meaningful and between cluster similarities are not guaranteed, we also plotted UMAP plots to better project the relationships of the individual neural subclusters (Fig. 6A, Fig. S7). SWNE analysis was also performed on the neural sub-cluster dataset to identify co-expressed genes (Fig. S8).

**Figure. 5:**
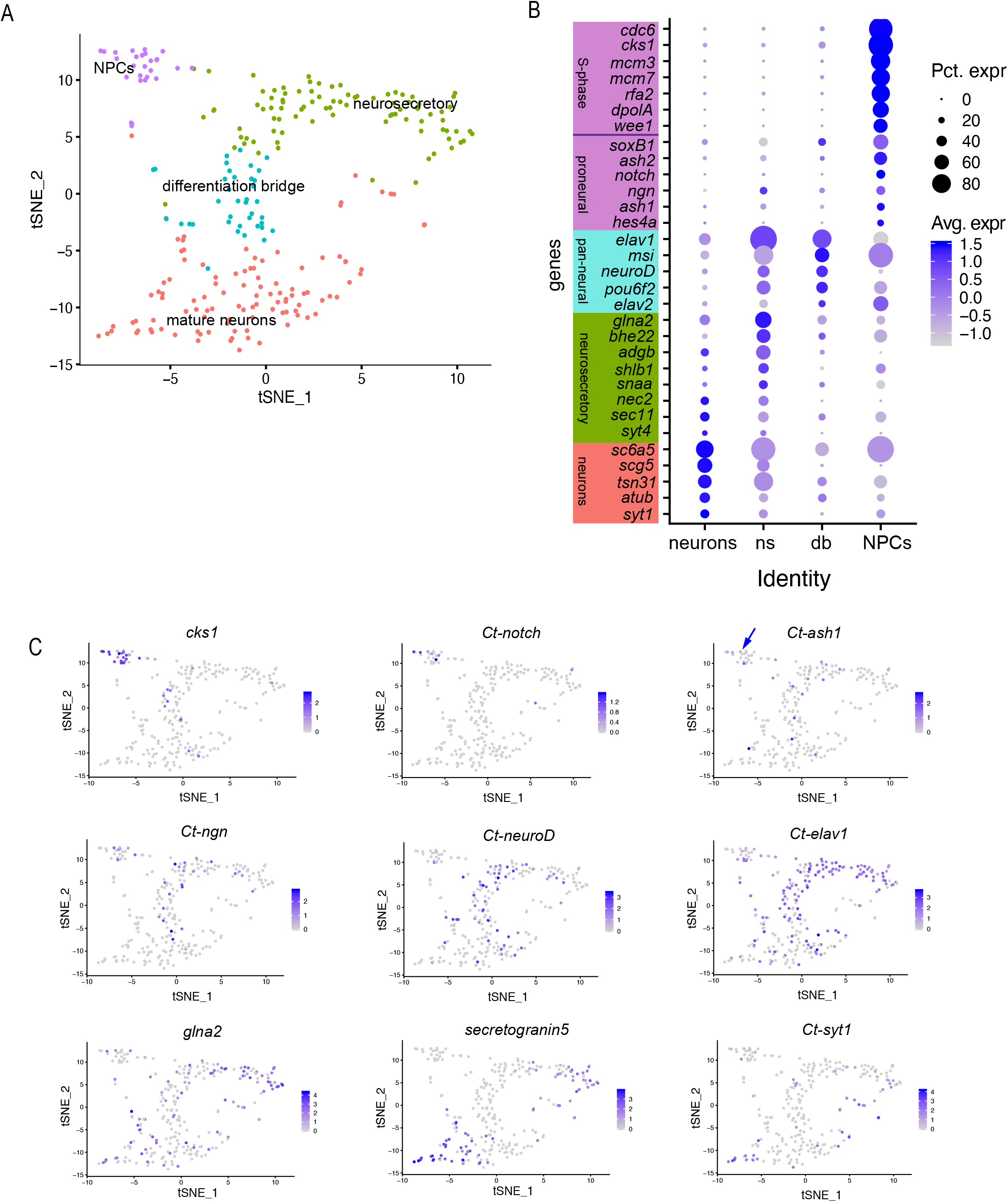
Neural cell type diversity in stage 5 larvae. (A) t-SNE representation of single cells obtained from the neural + neurosecretory cell clusters generated from unsupervised clustering of the stage 5 dataset, labeled and colored based on cluster identity (B) Dotplot showing differentially expressed genes per neural cell type identified. Each row represents a single gene regulating individual aspects of neurogenesis whereas each column represents one of the four neural cell types. The expression is log normalized per gene. (C) t-SNE plots colored by expression of selected marker genes that were used for identifying each cell type. The color key indicates expression levels (purple: high; grey: low). Blue arrow highlights expression of *Ct-ash1* in the NPC cluster. db, differentiation bridge; ns, neurosecretory.

**Figure 6:**
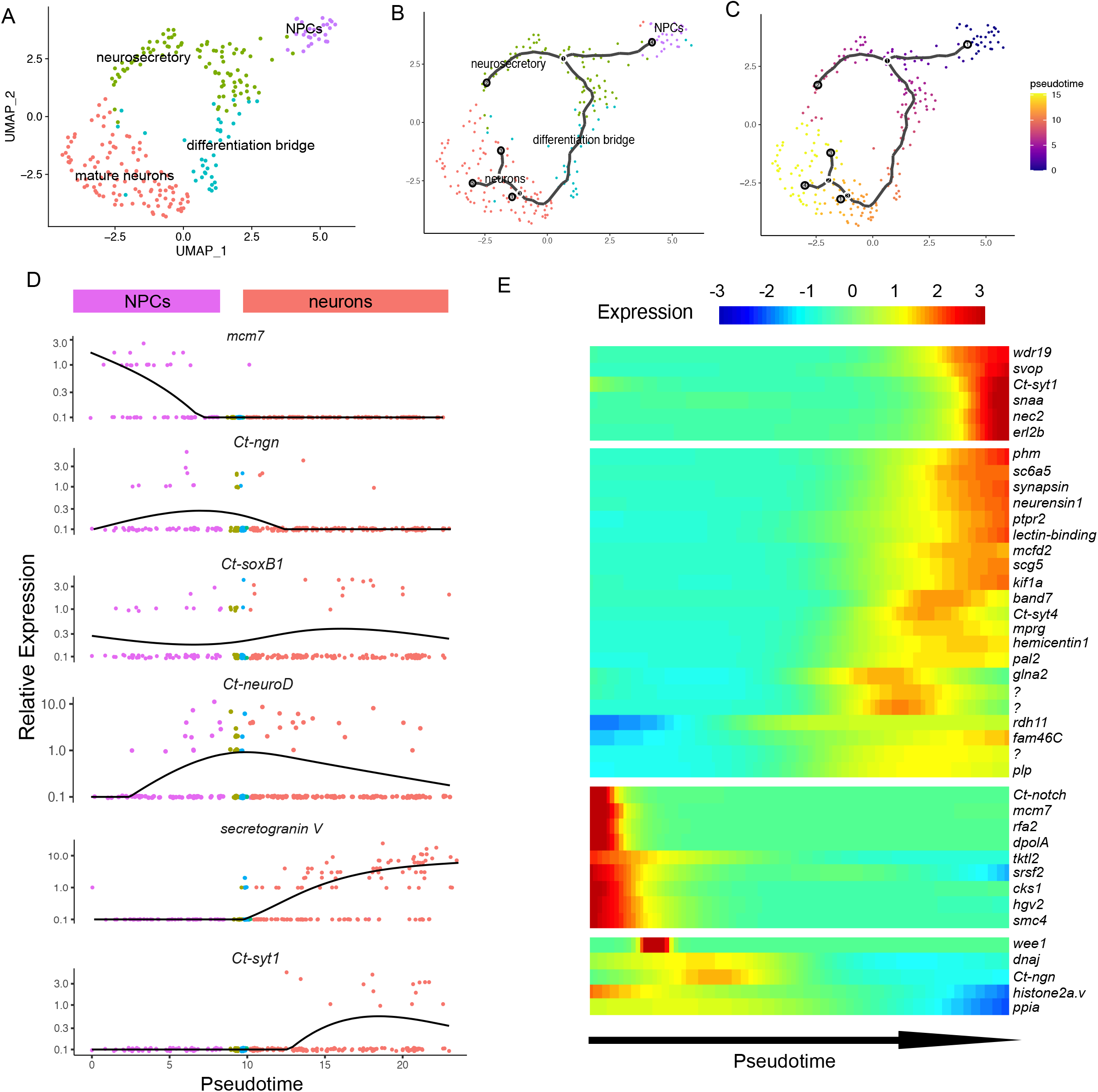
Lineage relationships between neural cell types and pseudotime analysis. (A) UMAP representation of single-cells from the neural + neurosecretory cell-cluster generated in the stage 5 unsupervised clustering. (B) Trajectory analysis using Monocle3 predicts a neural differentiation trajectory that begins with NPCs and ends with the mature neuronal and neuroendocrine cell cluster. The root of the trajectory lies within the NPC cell-cluster. (C) Monocle3 pseudo-temporal ordering of neural cells superimposed on the Seurat UMAP plot. Cells are colored based on their progression along pseudo-temporal space from pseudotime 0 in violet to the end of differentiation in yellow. (D) Monocle analysis predicts progressive expression dynamics of *mcm7* homolog, *Ct-ngn, Ct-soxB1, Ct-neuroD, secretogranin-V*, and *Ct-syt1*. (E) Heatmaps showing most significant TFs and effector genes clustered by pseudotemporal expression pattern (q values < 0.01). Pseudo-temporal ordering is from left (NPCs) to right (differentiated neurons). Selected transcription factors are shown for each cellular state along the differentiation trajectory.

Based on our t-SNE plot, we identified four clusters within the combined neural and neurosecretory group: (a) undifferentiated neural progenitors, (b) intermediate differentiation bridge, (c) differentiating neurosecretory cells and (d) mature neurons/neurosecretory cells containing a mixture of neurons with both neurotransmitter and neurohormonal output (Fig. 5A). The undifferentiated progenitors were identified based in the expression of S-phase markers such as *cdc6, cks1, rfa2, dpolA, wee1* and replication licensing factors such *mcm3* and *mcm7* (Fig. 5B, C) as well as M-phase markers such as *ccnb, mpip* and *cdk1*. These cells were also found to exclusively express *Ct-notch* (Fig. 5B, C). *Ct-notch* is expressed in both surface and subsurface cells the anterior neuroectoderm at stage 5 (Meyer and Seaver, 2009). We have previously shown in the *C. teleta* anterior neuroectoderm that surface cells primarily comprise rapidly dividing neural precursor cells (NPCs) while subsurface cells are largely post-mitotic neural cells (Meyer and Seaver, 2009; Sur et al., 2017; Sur et al., 2020). Hence this cluster may represent a combination of rapidly dividing NPCs and a few progenitors with limited proliferative potential. We named this cluster “NPCs”. Similar to our previous observations using EdU and fluorescent in-situ hybridization (FISH) (Sur et al., 2020), we observed *Ct-ash1* and *Ct-ngn* expression in this cluster (Fig. 5B, C, blue arrow). These undifferentiated cells were also found to express *Ct-msi* and *Ct-elav2* albeit at a much lower level (Fig. 5B) indicating that neural progenitors possibly express *Ct-msi* at lower expression levels.

We detected two transitional differentiation states (blue and green populations), one uniquely expressing pan-neural markers like *Ct-msi* and *Ct-elav2* that we named “differentiation bridge” and the other uniquely expressing *glutamine synthetase* (*glna2*), *androglobin* (*adgb*), *endophilin-1 (shlb1), neuroendocrine convertase* (*nec2*), and *synaptotagmin-4* (*syt4*) (Fig. 5A, B). *Glna2* is an enzyme involved in glutamine synthesis in excitatory glutaminergic neurons and has been shown to regulate the secretion of various adenohypophyseal hormones (Hrabovszky and Liposits, 2008). *Syt4* was found to be expressed in the neuroendocrine center of the vertebrate hypothalamus regulating oxytocin secretion (Zhang et al., 2011) as well as in the neuroendocrine center of the *P. dumerlii* head (Kaia Achim, 2017). Based on the expression of these genes, which are involved in the neuroendocrine pathway, we named this cell cluster (green population) “neurosecretory”. Both the intermediate differentiation bridge and differentiating neurosecretory cells (blue and green), expressed *Ct-elav1, Ct-neuroD* and *Ct-pou6* (Fig. 5B, C, Fig. 6A, Fig. S7A). A subset of cells in the “differentiation bridge” also expressed *Ct-ngn* and *Ct-ash1* indicating a later role in *C. teleta* neurogenesis (Fig. 5B, C).

The fourth cell population within the neural cluster expressed mature neuronal markers such as *synaptotagmin-1 (Ct-syt1), alpha-tubulin, Ct-synapsin* as well as neurosecretory markers such as *Ct-syt4* (Fig. 5B, C). Within the mature neuronal cell type, we detected a variety of neuronal subtypes: (i) glutaminergic neurons expressing *glutamine synthetase* (*glna2*) and *vesicular glutamate transporter* (*vgl2b*), (ii) cholinergic neurons expressing *acetylcholinesterase* (*aces*) and *vesicular acetylcholine transporter* (*vacht*), (iii) GABAergic neurons expressing *sodium- and chloride-dependent GABA transporter 1* (*sc6a1*), and (iv) neuroendocrine subtypes expressing *Ct-syt4, secretogranin-V* and *prohormone-4* among others (Fig. 5A–C). A subset of cells in the “mature neuron” cluster expressed these neuroendocrine genes at a much higher level than observed in the differentiating neurosecretory cells (green population), possibly indicating that these are mature neuroendocrine cells that clustered with the other mature neuronal subtypes (Fig. 5B). Overall, our t-SNE and UMAP analyses highlighted different neural cell types that were previously reported (Meyer and Seaver, 2009; Sur et al., 2017; Sur et al., 2020) as well as previously unknown neuronal subtypes within each cell-cluster that await further characterization.

Next, we applied SWNE analysis to identify coregulated gene modules within each neural cell type (Fig. S8). We observed different sets of co-expressed genes in the undifferentiated cluster. One such coregulated subset of genes included *cks1* (cyclin-dependent kinase regulatory subunit 1), *bafB* (Barrier to autointegration factor B) and *hgv2* nucleosomal assembly factor. All three genes in this module play important roles in cell-cycle progression (Furukawa et al., 2003) (Fig. S8A, B). The differentiation transition states were also found to co-express genes such as *neuroD, dpys* and *rdh11* and *talin-1* indicating that cells in this cluster are already on distinct neural differentiation trajectories. *Talin-1* was expressed in another gene module present in the differentiating neurosecretory cells along with an EF-hand domain containing protein and a gene encoding a potassium voltage-gated channel subfamily H8 (*kcnh8*). Among the cells that clustered within the mature neuronal cluster, we detected subsets of cells expressing genes encoding V-type proton ATPase (*vatl*) as well as peptidergic neuronal markers such as *myom1* and *orckB* (Fig. S8A, B).

### Computational lineage reconstruction reveals temporal relationships between neural cell types

To understand pseudotemporal relationships between the different neural cell types at stage 5, we used Monocle3.0.2, which orders cells based on similarities of their global transcriptional profiles. Starting from the neighborhood graph generated in t-SNE or UMAP space (Fig. 6A), Monocle uses reversed graph embedding to reconstruct single-cell trajectories in a fully unsupervised manner (Trapnell et al., 2014a; Qiu et al., 2017). Using Monocle, we also identified variable gene sets or modules in different cell states (Fig. S9). While running the Monocle3 algorithm without any assumptions about the trajectory, we obtained an abstracted graph that allowed us to derive a single differentiation tree that included all the neural cell types and linked them to one root, the NPC cluster (Fig. 6B). Along the trajectory, cells were ordered based on their developmental origin and state of differentiation (Fig. 6A, B). This generated a pseudotime trajectory with six distinct cell states (Fig. 6C–E). These were defined by the expression of *Ct-notch* and S-phase markers (*mcm7, rfa2, dpolA, cks1*) for the NPC state; *Ct-ngn* and *wee1* for a progenitor state; *kif1a, band7, mprg, hemicentin-1*, and *Ct-syt4* for an intermediate neuronal differentiation state; *pal2, glna2, plp*, and *rdh11* for an intermediate neurosecretory differentiation state; *Ct-syt1, Ct-synapsin, synaptobrevin, neurensin, neuroendocrine convertase-2* (*nec2*) among others in the final state comprising both mature neuronal and neuroendocrine cell types (Fig. 6E, Fig. S7A–G). Interestingly, the proximity of these cell types in the UMAP plot (Fig. 6A) indicated that their transcriptomes are closely related in a continuous fashion.

To identify temporal progression of genes that may be involved in neurogenic cell fate decisions, we mapped some previously characterized genes that significantly varied in their pseudotemporal expression and looked more closely at their expression dynamics (Fig. 6C). This analysis showed several discrete shifts in gene expression patterns during *C. teleta* neurogenesis. For example, S-phase markers (*mcm7, rfa2* and *dpolA*) and *Ct-notch* were only expressed in proliferating NPCs and were rapidly downregulated at the onset of differentiation (Fig. 6D, E). In another subset of NPCs (Fig. 6A–C), genes like *wee1, Ct-ngn* and other bHLH transcription factors such as *Ct-ash1* and *Ct-atonal* were upregulated later in pseudotime than the S-phase markers and were not downregulated until the latter stages of neural differentiation (Fig. 6D, E; Fig. S9). Expression of *Ct-neuroD* peaked as *Ct-soxB1* and *Ct-ngn* began to become downregulated (Fig. 6D). Such an observation closely follows patterns obtained using double-FISH and FISH+EdU experiments reported previously (Sur et al., 2020). Genes involved in imparting a neurosecretory identity such as *secretogranin-V, pal2* and *glutamine synthestase* (*glna2*) were found in the next step of the cascade of differentiating cells (Fig. 6A–E). These genes turned on as *Ct-neuroD* expression began to decline (Fig. 6D) but were downregulated prior to the expression of the next subset of markers such as *Ct-syt1, Ct-syt4, Ct-synapsin, neurensin*, and *synaptobrevin* (*snaa*) among others, which initiated their expression and increased later in pseudotime (Fig. 6E, Fig. S7G). These late-expressing genes like *Ct-syt1, Ct-syt4, Ct-synapsin* and *snaa* likely modulate neurotransmitter and neurohormonal release in the presynaptic cleft and hence are expressed in mature neurons. Therefore, our pseudotemporal analysis revealed the onset of the neurosecretory program prior to the neuronal program. Neural subtype specific markers such as *acetylcholinesterase* (*aces*), *vesicular acetylcholine transporter (vacht), glna2* and *sc6a1* were expressed even later in pseudotime and represent different neuronal subtypes such as cholinergic and GABAergic neurons (Fig. 6E, Fig. S7D). Interestingly, we also observed expression of *neuroendocrine convertase* (*nec2*) in this pseudotemporal cluster indicating some mature neuroendocrine cells as well. Overall our pseudotemporal analysis elucidated two differentiation trajectories from undifferentiated progenitors to mature neuronal or neurosecretory cell types and identified both previously-known and unknown markers for neurogenesis along both trajectories in *C. teleta*.

### RNA velocity analysis confirms lineage relationships predicted by Monocle3

To independently validate the differentiation trajectories predicted by Monocle3 and to gain insight into dynamics of stem-cell activation and differentiation, we used velocyto (La Manno et al., 2018), a computational method that tracks recent changes in transcriptional rate of a gene to predict future mRNA levels of that gene (Fig. 7A, B). These transcriptional rate changes are estimated for each gene by calculating the ratio of spliced versus unspliced reads in the sequencing data (Fig. S10A, B; Fig. S11) and extrapolating over all genes across all cells in the dataset. The timescale of future cell-state prediction is on the scale of a few hours (La Manno et al., 2018).

**Figure 7:**
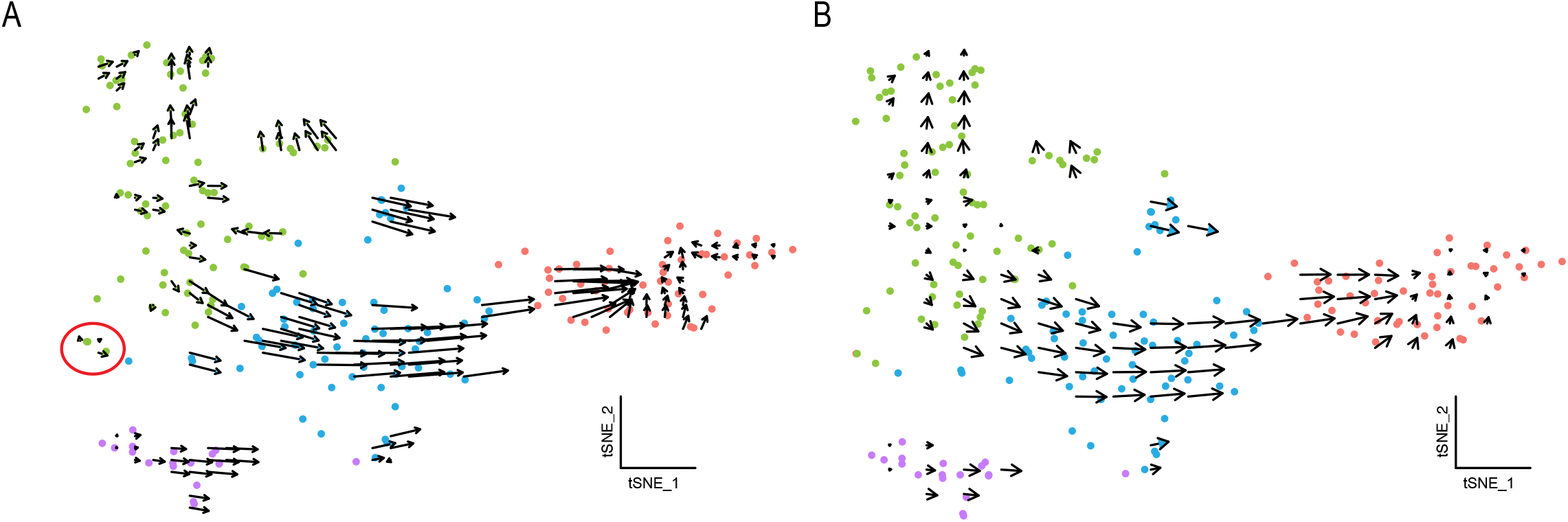
RNA velocity plotted in t-SNE space for neural cell types. (A) Velocyto force field showing the average differentiation trajectories (velocity) for cells located in different parts of the tSNE plot. For each cell, arrows indicate the location of the future cell state. RNA dynamics vary between NPCs, the two differentiation trajectories and mature neurons and also within each cell type. Velocity estimates based on nearest cell pooling (k=5) were used. Red circle shows two cells with velocity fields pointing along the two differentiation trajectories possibly representing a progenitor population for neural and neurosecretory cells respectively. (B) Same velocity field as A, visualized using Gaussian smoothing on a regular grid.

We estimated RNA velocity for each cell within the combined neural and neurosecretory group at stage 5 (C3 + C4) to assess the relationship between NPCs, differentiating neurons and mature neurons. We projected the estimated cell states onto the t-SNE plot, which describes the path predicted by the RNA velocity algorithm and visualized the results by plotting an arrow for each cell spanning its actual and predicted future cell state. Hence, cells that are transcriptionally active have long arrows, whereas cells that are undergoing very low transcriptional turnover have either short or no arrows. For example, in the mature neuronal cell-cluster we observed little and uncoordinated RNA velocity indicating that these cells are transcriptionally stable and are undergoing less changes at the RNA level, reinforcing that these cells represent terminally differentiated cell types (Fig. 7A, B). Projecting the RNA velocity of individual cells states on a PCA plot separated each cell cluster and captured the main neural differentiation axis (Fig. S10C–F). Similarly, within the NPC cluster (purple), a subset of cells exhibited very short arrows indicating very low RNA metabolism whereas another subset was found to have relatively longer arrows pointing along the differentiation axis (Fig. 7A, B). Finding differential transcriptional activity in this cluster highlights that this is a heterogenous population, similar to what we detected using Monocle3 pseudotemporal analysis (Fig. 6E).

We observed differences in RNA velocity between the two differentiation trajectories as well. In the neuronal differentiation trajectory, RNA velocity was higher than that in the neurosecretory trajectory. However, arrows within both trajectories pointed away from the NPCs indicating the direction of differentiation. Decreasing RNA velocity towards the far end of the neurosecretory trajectory led us to speculate that these cells have a stable transcriptome and may represent terminally differentiated neurosecretory cells. We identified the root of the differentiation process in the NPC cluster similar to our observations from pseudotime analysis and trajectory prediction using Monocle3. We also identified two cells that lie close to each other in t-SNE space and point along the direction of each differentiation trajectory, possibly representing neural and neurosecretory progenitors, respectively (Fig. 7A, red circle). The paths predicted by velocyto largely agree with the trajectories predicted by Monocle, which supports two independent differentiation trajectories for neurons and neurosecretory cells. The continuity of cells in t-SNE space obtained using our RNA velocity analysis (Fig. 7A, B) coupled with pseudotime analysis (Fig. 6C–E) and gene expression across cell types (Fig. 5B, C) highlights the continuous nature of *C. teleta* neurogenesis, making rigid classification of neural cell types along the differentiation trajectory difficult.

## Discussion

We present here one of the first scRNAseq studies on a spiralian annelid using the 10X genomics platform. Our experimental design and analysis revealed that 10X genomics may not be the optimal platform for sequencing dissociated cells from spiralian larvae due to their smaller cell size as compared to most well-studied vertebrate taxa. Due to the automated droplet-generation and cell-capture procedures as part of the 10X workflow, we suspect that our cells may not have been captured efficiently and that resulted in low RNA recovery. Therefore, future scRNAseq approaches in spiralian taxa need to be designed using manual capture of cells more along the lines of SmartSeq approaches that have the potential to generate high-quality libraries from low RNA input samples (Satija et al., 2015; Farrell et al., 2018). However, SmartSeq approaches are less cost-effective than bulk droplet-capture procedures like 10X genomics and DropSeq (Ziegenhain et al., 2017), making future efforts at optimizing these techniques for spiralians important.

Despite the technical challenges of using 10X genomics with cells of small sizes, our study allowed us to investigate gene expression, developmental trajectories and identify molecular domains in larvae at two different stages for the annelid *Capitella teleta*. Using this approach, we demonstrate that (i) previously characterized marker genes in *C. teleta* can be used to annotate bioinformatically derived cell-clusters, (ii) Louvain clustering analysis can resolve the entire *C. teleta* body plan into distinct molecular domains based on differential gene expression, and (iii) trajectory analysis can track continuous changes in pseudotemporal gene expression patterns during differentiation of certain cell types. Recent work has only examined gene expression dynamics in *C. teleta* using WMISH and immunolabeling experiments, and we view our study as a first step towards understanding annelid and eventually spiralian development at a systems level. Furthermore, we present the differential gene expression analyses and cell types in this study as hypotheses that require validation via functional and in situ expression studies to test the accuracy of the *in-silico* predictions made here.

### Molecular subdivision of the *C. teleta* larval body

Our data allow comparison of cell types at the single-cell level across the entire animal. Our unsupervised clustering approaches classified the *C. teleta* larval whole-body into eight distinct molecular domains – (i) generic precursor cells, (ii) ectodermal precursors, (iii) myoblasts, (iv) ciliary-bands, (v) gut secretory cells, (vi) neural cells, (vii) neurosecretory cells, and (viii) protonephridia. By comparing our findings with previous studies, we created an annotated list of markers for each of the cell clusters that were previously characterized as well as novel markers as highlighted in the Results section. Our multifaceted analyses revealed the progressive origin of tissues and how the genes underlying the development of these tissues change across the two larval stages. The first few discrete clusters to originate soon after gastrulation included gut, ciliary-band and nervous system. After 24 hours of development, more discrete clusters were identified including neurosecretory cells and protonephridia. Interestingly, neurosecretory properties were detected as early as stage 4 (~24 hours post gastrulation), but these cells clustered together with neurons. Based on previous work, neurons in *C. teleta* are first detected in the developing brain at stage 4 (Meyer and Seaver, 2009; Meyer et al., 2015; Sur et al., 2017). The presence of neurosecretory cells together with neurons in our stage 4 t-SNE plot indicates a considerable neuronal diversity at the initial stages of *C. teleta* development. Such early appearance of diverse neurons may enable *C. teleta* larvae to respond to environmental stimuli. Furthermore, from our pseudotemporal analysis on the neural and neurosecretory subclusters, we inferred progressive changes in gene expression during the progression of NPCs to neurons. Our data suggest a cascade in which early cell-cycle markers are rapidly downregulated followed by an upregulation of neural differentiation markers such as *Ct-neuroD* and subsequently by orthologs of genes involved in neuronal migration and terminal differentiation genes that function in mature neurons. Using a marker-independent approach, velocyto, we also computed RNA velocity within the neural cell clusters and recovered similar differentiation trajectories and transcriptional dynamics that we deduced using Monocle, showing the robustness of our approaches.

### Evolutionary comparisons of the *C. teleta* cell types with other annelids

Our *C. teleta* single-cell analysis presented here enables a systematic comparison of cell types across species. We found that cells at stages 4 and 5 in *C. teleta* expressed many genes shared with homologs in *P. dumerilii* larvae around a similar developmental stage (Kaia Achim 2017). For example, gut primordial cells of endodermal origin expressing *hnf4a* and *collagen alpha 3*, ciliary-bands expressing *rsph3, tekt4a, tekt1a*, and *ift80*, and myoblasts expressing *myosin* were found to be similar to that in *P. dumerlii* (Kaia Achim 2017). We observed consistent expression of *tektin* homologs in the ciliary band cluster across the two stages in *C. teleta*, and these homologs were previously reported in ciliary bands of *P. dumerlii* and another annelid *Hydroides elegans* (Arenas-Mena et al., 2007; Kaia Achim, 2017). Recent reports show the expression of *tektin* homologs in the *P. dumerilii* prototroch, ciliated apical organ, telotroch and two pairs of paratrochs, and axonemal dyenin homologs in all ciliary structures of the mollusk *Tritia obsoleta* (Wu et al., 2020b), similar to our findings in *C. teleta*. This suggests that a role of *tektin* homologs in the ciliary bands of annelid larval trocophores may have been a conserved feature. A recent report by Wu et al., 2020b uncovered two spiralian-specific genes expressed in the ciliary bands of most spiralians called *lophotrochin* and *trochin*. These genes along with the markers identified in our study provide a valuable resource in further characterization of the origin of ciliary bands within Spiralia.

Some of gut cells described here express *Ct-blimp1* and represent endodermal midgut precursors (Boyle et al., 2014). In both *C. teleta* and *P. dumerilii*, the large, yolky midgut cells originate from the vegetal macromeres (Ackermann et al., 2005; Meyer et al., 2010). We only noticed shared expression of *hnf4a* between both annelids but not expression of any of the other *P.dumerlii* “gut” markers in our dataset (Kaia Achim, 2017). We think it is likely that we may have size-excluded a majority of large, yolky midgut cells at stages 4 and 5 that were larger than 40 μm in size prior to cell-capture (see Materials and methods). Whether the genetic developmental program is conserved between the two annelids needs more investigation.

Within the neural and neurosecretory cell types, we also identified markers that were previously detected in the scRNAseq dataset for *P. dumerlii* larvae (Kaia Achim 2017). Subclustering these cell types allowed us to detect coherent sets of effector genes and transcription factors expressed at different pseudotimes, representing distinct cellular modules, e.g. NPCs, intermediate differentiation bridge, differentiating neurosecretory cells, and mature neurons and neuroendocrine cells. In addition, we also observed considerable differences in gene expression even within individual neural subgroups highlighting distinct but related cell types. For example, within the NPC cluster, we observed two different gene modules that were expressed at different pseudotimes and had differential transcriptional activity, indicating a heterogenous population of NPCs. From our subclustering analysis of neural cells, we identified putative *phc2^+^* neurosecretory cells in *C. teleta*, which may be homologous to *phc2*^+^ neurosecretory centers in *P. dumerlii* and other spiralians (Tessmar-Raible et al., 2007; Kaia Achim, 2017). Phc2^+^ neuroendocrine centers were also detected apically in the developing larval brain of *P. dumerlii* are were found to express other vertebrate-type neuropeptides such as the Vasotocin/neurophysin preprohormone (Kaia Achim, 2017). Vasotocin/neurophysin homologs have been found in many spiralians including annelids (Oumi et al., 1994; Tessmar-Raible et al., 2007), gastropods (van Kesteren et al., 1992), and cephalopods (Takuwa-Kuroda et al., 2003). However, we could not detect a vasotocin/neurophysin homolog in *C. teleta* from our scRNAseq analysis although we only detected one or two conopressin/neurophysin-expressing cells at stages 4 and 5. Therefore, presence of larval neuroendocrine centers regulating neurohypophyseal hormonal activity seems to be a conserved feature among spiralians that has been lost in *D. melanogaster* and *C. elegans* (Tessmar-Raible et al. 2007).

Altogether, our *C. teleta* scRNAseq study suggest that comparative studies of neural cell types across animal evolution using high-throughput scRNAseq is a promising direction for evo-devo research and needs to be expanded to more taxa. As exemplified here, whole-organism scRNAseq across many taxa can provide comprehensive insights into metazoan cell type evolution and tissue-specific genome-wide regulatory networks.

## Author Contributions

Conceptualization: A.S., N.P.M.; Methodology: A.S., N.P.M.; Software: A.S.; Formal analysis: A.S.; Investigation: A.S.; Writing - original draft: A.S., N.P.M.; Writing - review & editing: A.S., N.P.M.; Supervision: N.P.M; Funding acquisition: A.S., N.P.M.

## Acknowledgements

The authors are grateful to the Boston University Single-Cell Sequencing Core for their services and their excellent support.

