## Supplementary Figures and Tables for "Resolving transcriptional states and predicting lineages in the annelid *Capitella teleta* using single-cell RNAseq"

### Supplementary Material

#### Supplementary Figures

A

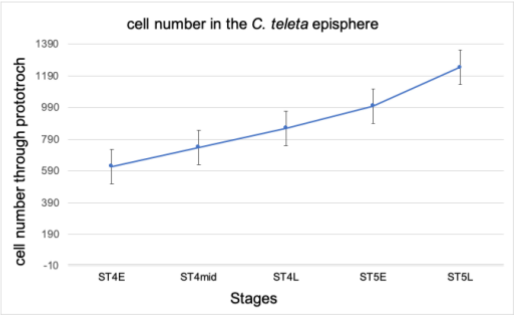


B

| Stage | Number of animals scored | Mean number of cells in the episphere (± S.E.M.) |
|---|---|---|
| ST4 early | 10 | 617.1 ± 14.756 |
| ST4 mid | 8 | 738.5 ± 15.578 |
| ST4 late | 7 | 859.4 ± 21.864 |
| ST5 early | 6 | 997.8 ± 30.040 |
| ST5 late | 8 | 1242.1 ± 77.932 |

C

| Enzyme | Concentration | No. of replicates | Total cells Recovered (n=300) (± S.E.M) | % of cells recovered per larvae (± S.E.M) | Proportion cell survival (± S.D.) |
|---|---|---|---|---|---|
| Papain | 1% | 5 | 119166 ± 27577 | 9.930 ± 2.298 | 0.844 ± 0.017 |
| Papain | 2% | 3 | 123333 ± 20989 | 10.277 ± 1.749 | 0.739 ± 0.007 |
| Trypsin | 1% | 6 | 105000 ± 17819 | 8.750 ± 1.484 | 0.965 ± 0.003 |
| Trypsin | 2% | 3 | 93300 ± 17766 | 7.775 ± 1.480 | 0.960 ± 0.002 |
| Papain + Trypsin | 1% | 3 | 99165 ± 30007 | 8.263 ± 2.500 | 0.857 ± 0.008 |
| Pronase | 1% | 3 | 95000 ± 10832 | 7.916 ± 0.902 | 0.916 ± 0.021 |

D

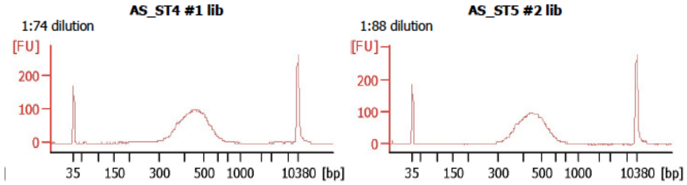


E

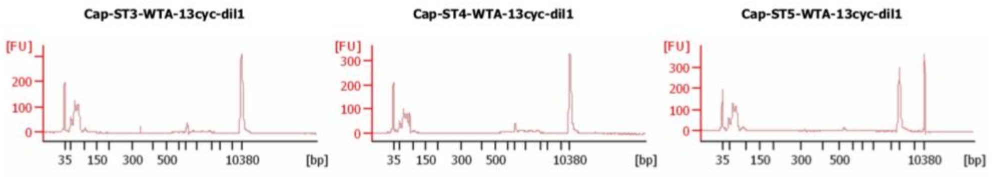

**Figure S1: Quantification of total number of cells in the *C. teleta* developing head and optimizing cell dissociation protocols.** (A) Line graph showing the total number of Hoescht$^+$ cells counted from the apical surface of the head through the prototroch across stages 4 to 5 using confocal z-stacks. X-axis shows the different stages quantified and the y-axis shows the total number of cells counted. (B) Table showing the average number of cells (± S.E.M) in the head across stages 4 to 5 quantified from confocal z-stacks. Number of animals quantified for each stage shown in the second column. (C) Table showing the different proteolytic enzymes tested across different concentrations for optimizing *C. teleta* cell dissociation protocols using stage 4 larvae. The first two columns indicate the enzymes and the respective concentrations used. The third column shows the number of biological replicates for each enzyme conducted, the fourth column indicates the total number of cells (± S.E.M) quantified post-dissociation from 300 stage 4 larvae, the fifth column indicates the percentage of cells recovered per stage 4 larvae (± S.E.M) and the sixth column shows the proportion of healthy cells (± S.D.) quantified following a Trypan Blue exclusion test 1 h post-dissociation using the respective enzymes. (D) Bioanalyzer data showing quality of the 10X 3'v3 sequencing libraries from our first trial at Boston University. (E) Bioanalyzer data showing no evidence of cDNA traces following whole transcriptome amplification after 13 cycles of amplification from 4000 cells per stage (stage 3, stage 4 and stage 5) during our second trial at Harvard University. ST4E, stage 4 early; ST4mid, stage 4 middle; ST4L, stage 4 late, ST5E, stage 5 early; ST5L, stage 5 late.

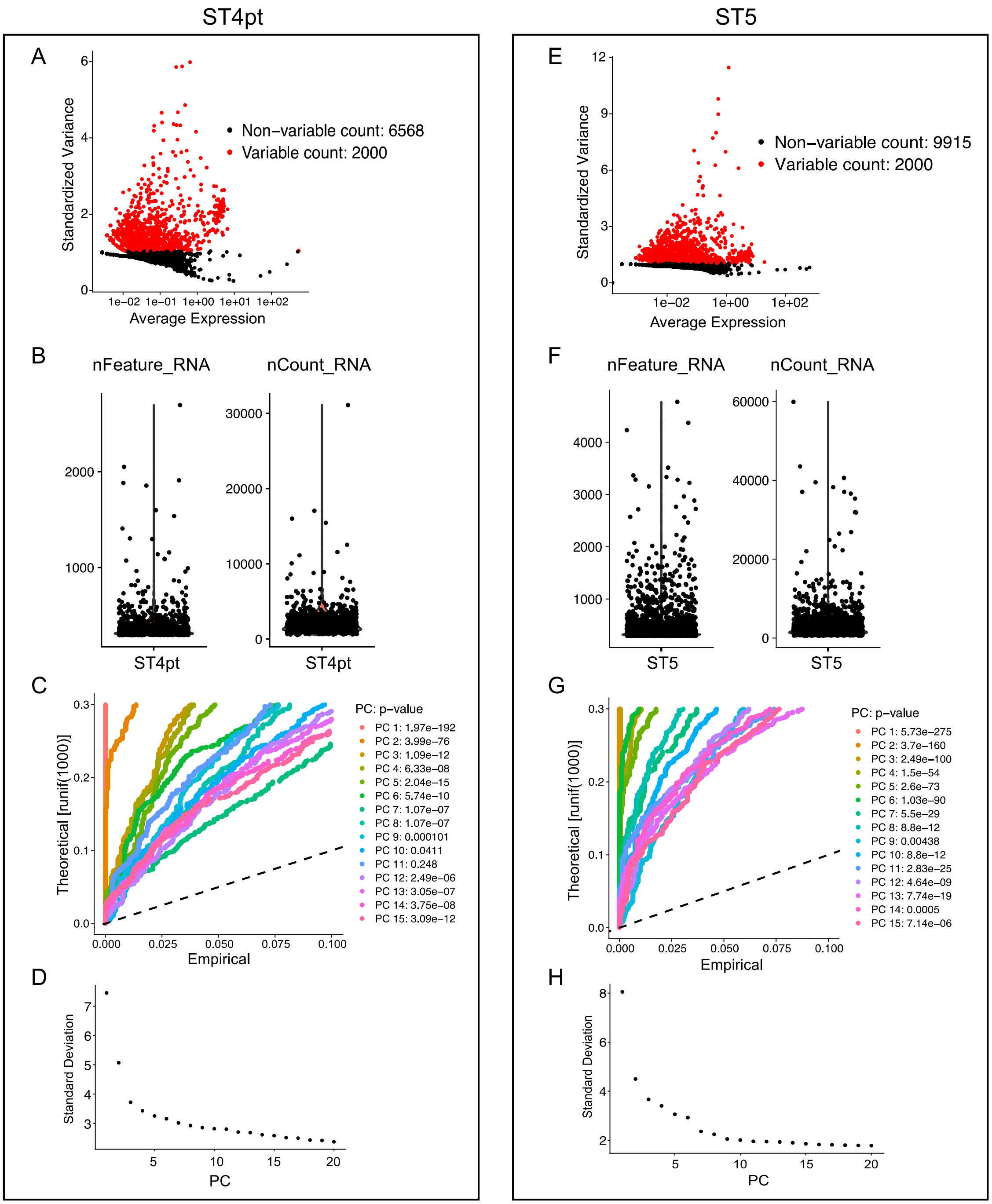


**Figure S2: Statistics on generating high-quality single-cell transcriptomes at stage 4 and 5.** (A, B) Top 2000 most variably expressed genes at stage 4 (A) and stage 5 (B) were identified and used for further downstream identification of significant principal components. (C, D) Plots showing the feature selection for both stage 4 (C) and stage 5 (D) datasets. Cells were selected such that they had fewer than 2500 features to exclude cell-doublets. (E, F) Jack-Straw plots of principal components for stage 4 (E) and 5 (F). (G, H) Elbow plots at stages 4 (G) and (H) showing standard deviations of the principal components.

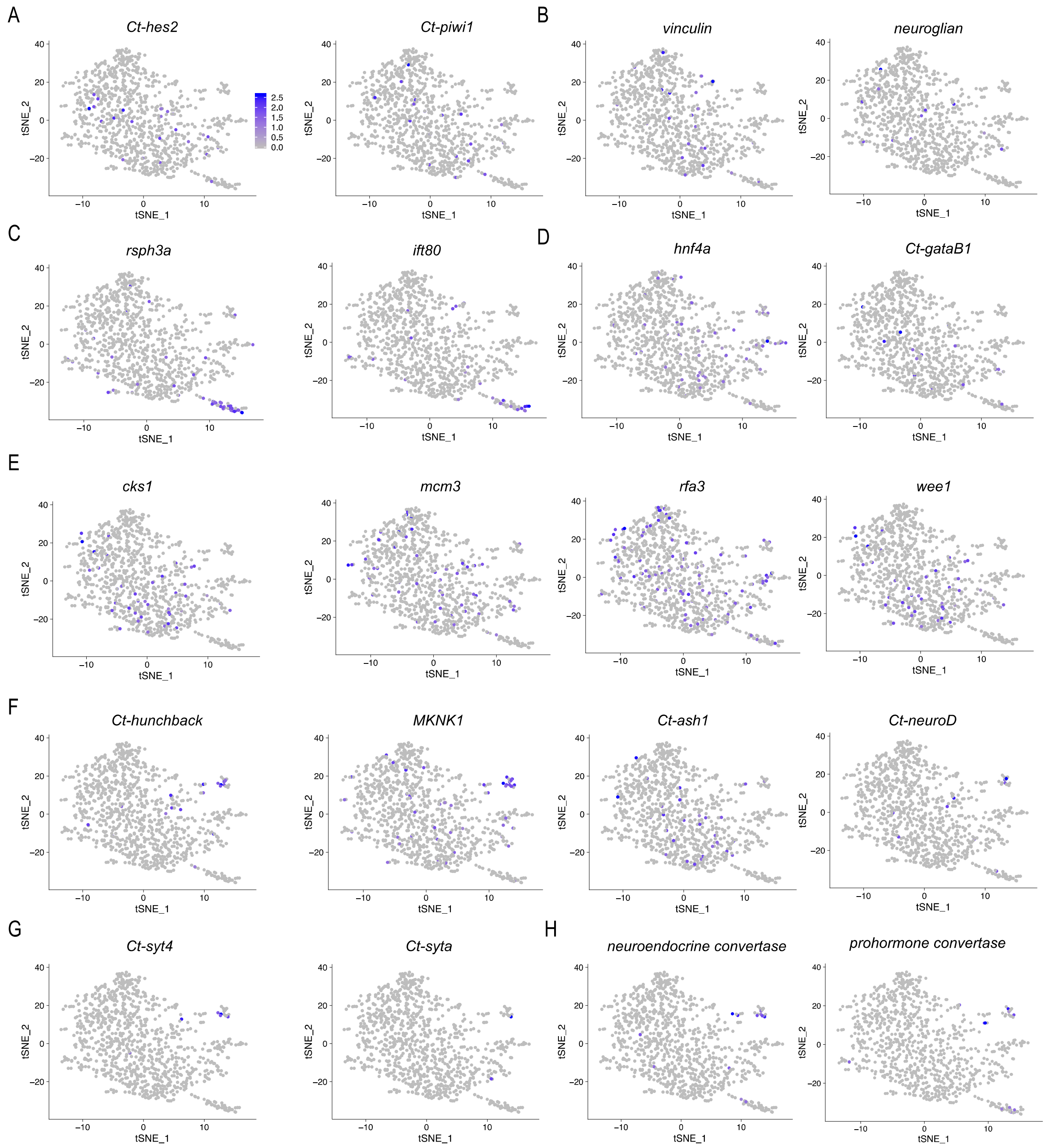


**Figure S3: Cluster identities of stage 4 dataset.** For annotated clusters, see Fig. 3A. (A) precursors – *Ct-hes2*, *Ct-piwi1*. (B) ectodermal precursors – *vinculin*, *N-cadherin*. (C) ciliary-band – *Rsph3a*, *tektin1a*. (D) gut digestive enzymes – *hnf4a*, *Ct-gataB1*. (E) S-phase and M-phase markers – *cks1*, *mcm3*, *rfa3*, *wee1*. (F) Neural markers and bHLH transcription factor homologs – *Ct-hunchback*, *MKNK1*, *Ct-ash1* and POU homeodomain factor *pou6f2*. (G, H) mature neuronal and neurosecretory markers such as *Ct-syt4*, *Ct-syta* (G) and neuroendocrine markers such as *neuroendocrine convertase*, *prohormone convertase* (H).

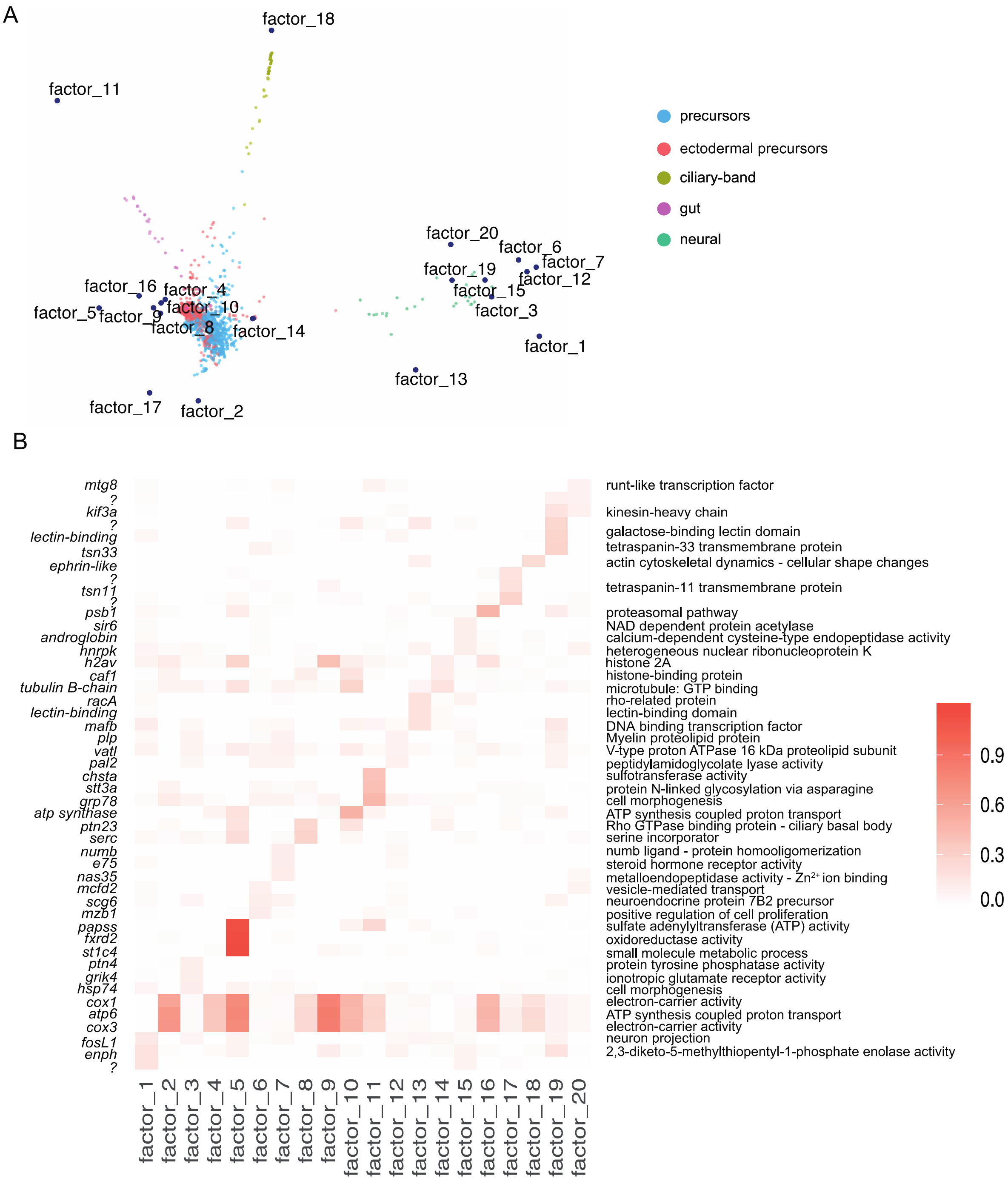


**Figure S4: SWNE visualizations to capture global and local structure in the stage 4 larval dataset.** (A) SWNE (k = 20) representation of stage 4 larval single cells with clusters labeled by molecular identities and gene modules represented as factors embedded within the SWNE visualization. (B) Heatmap showing the top factors ($p < 0.01$) identified with SWNE analysis using GO-term based gene sets that vary across all cells. The GO terms associated to each factor are shown on the right. Each row represents a gene and each column represents a factor expressed in a particular cluster based on the SWNE embeddings shown in A. The clustering obtained with SWNE defines groups of cells that strongly agree with Seurat clustering.

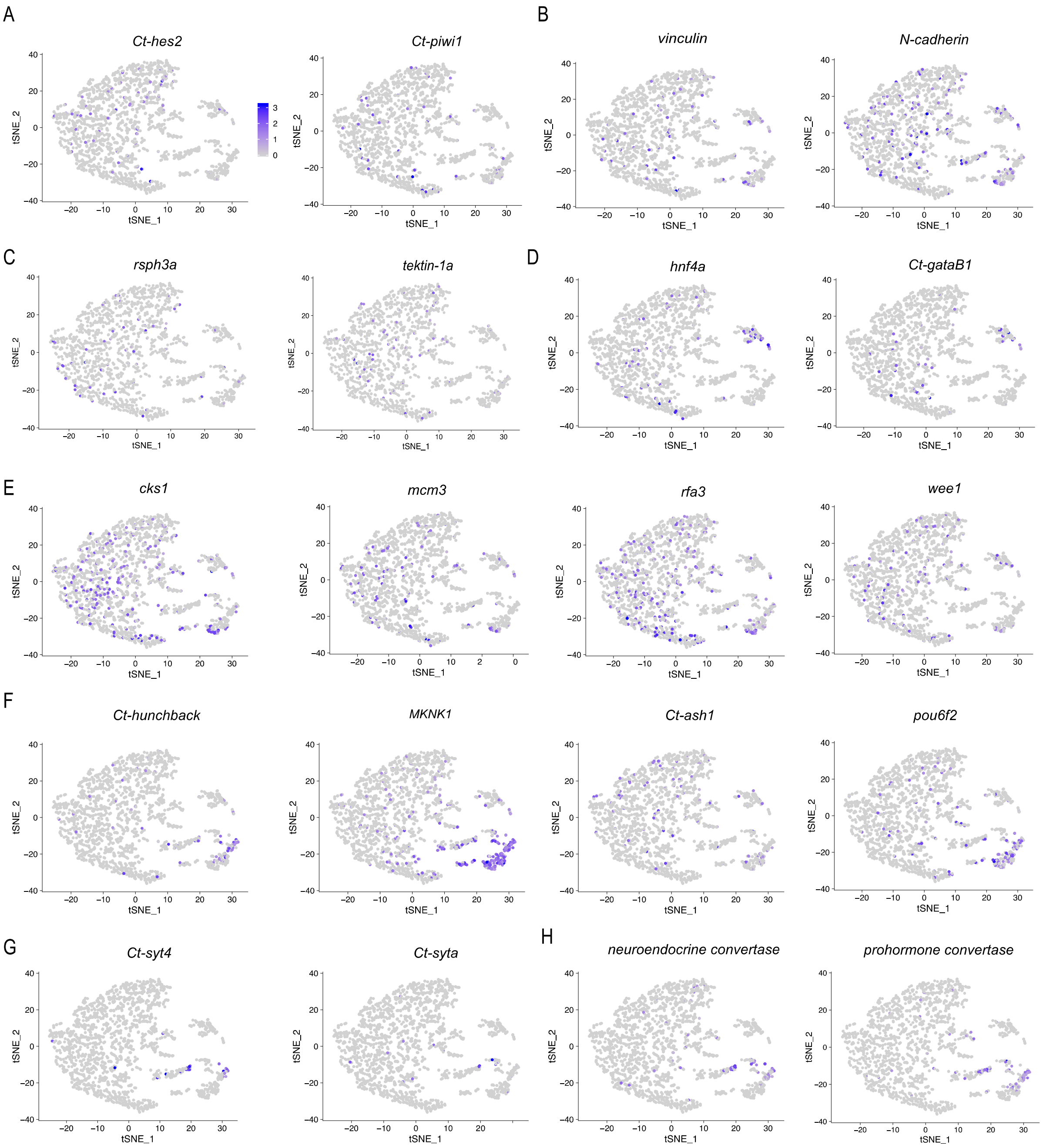


**Figure S5: Cluster annotation of stage 5 dataset.** For annotated clusters, see Fig. 4A. (A) precursors – *Ct-hes2*, *Ct-piwi1*. (B) ectodermal precursors – *vinculin*, *N-cadherin*. (C) ciliary-band – *Rsph3a*, *tektin1a*. (D) gut digestive enzymes – *hnf4a*, *Ct-gataB1*. (E) S-phase and M-phase markers – *cks1*, *mcm3*, *rfa3*, *wee1*. (F) Neural markers and bHLH transcription factor homologs – *Ct-hunchback*, *MKNK1*, *Ct-ash1* and POU homeodomain factor *pou6f2*. (G, H) mature neuronal and neurosecretory markers such as *Ct-syt4*, *Ct-syta* (G) and neuroendocrine markers such as *neuroendocrine convertase*, *prohormone convertase* (H).

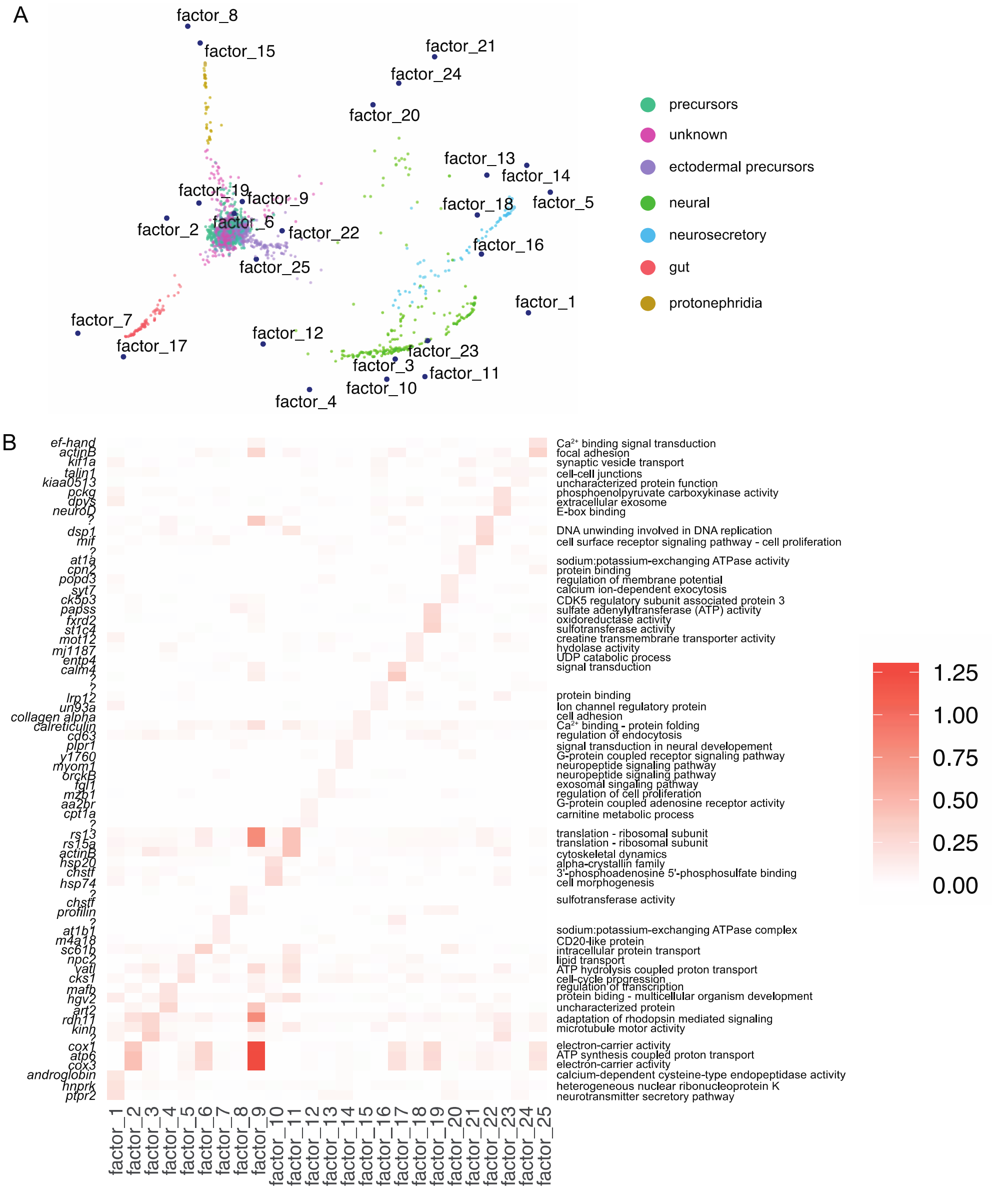


**Figure S6: SWNE visualizations to capture global and local structure in the stage 5 larval dataset.** (A) SWNE (k = 25) representation of stage 5 larval single cells with clusters labeled by molecular identities and gene modules represented as factors embedded within the SWNE visualization. Two more molecular domains detected in the stage 5 SWNE plot in comparison to stage 4. The protonephridia arise from the central ectodermal cluster while the neurosecretory cells branch off from the neural cell cluster. Most number of factor embeddings were observed in the neural and neurosecretory clusters indicating a high diversity in cell types expressing different sets of coregulatory genes. (B) Heatmap showing the top factors ($p < 0.01$) identified with SWNE analysis using GO-term based gene sets that vary across all cells. The GO terms associated to each factor are shown on the right. Each row represents a gene and each column represents a factor expressed in a particular cell type based on the SWNE embeddings shown in A.

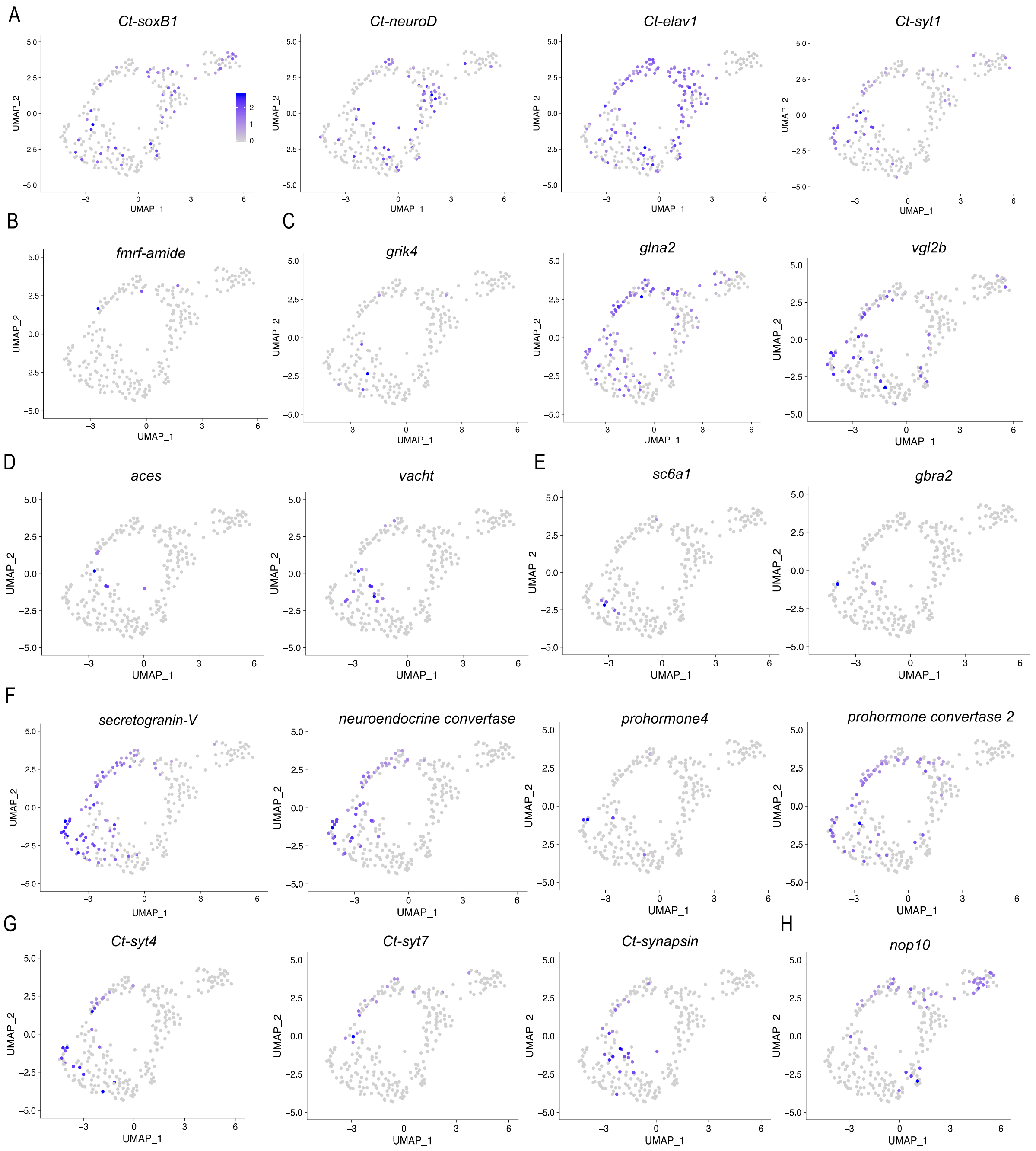


**Figure S7: Cluster annotation of stage 5 neural + neurosecretory subclustered dataset (Related to Fig. 6 and S5).** For annotated clusters, see Fig. 5A. (A) Previously characterized neurogenic homologs – *Ct-soxB1*, *Ct-neuroD*, *Ct-elav1*, *Ct-syt1*. (B) RFamide$^+$ neurons – *fmrfamide* (C) glutaminergic neurons and glutamate receptors – *grik4*, *glna2, vgl2b*. (D) cholinergic neurons – *aces*, *vacht*. (E) GABAergic neurons – *sc6a1*, *gbra2*. (F) neuroendocrine precursors – *scg5*, *nec2*, *prohormone4*, *prohormone convertase 2 (phc2)*. (G) Ca2$^+$ independent syntaptotagmins – *Ct-syt4*, *Ct-syt7* (H) snoRNA binding proteins – *nop10*. The expression bar in (A) indicates upregulation and downregulation of genes in each feature-plot (gray: downregulated, purple: upregulated).

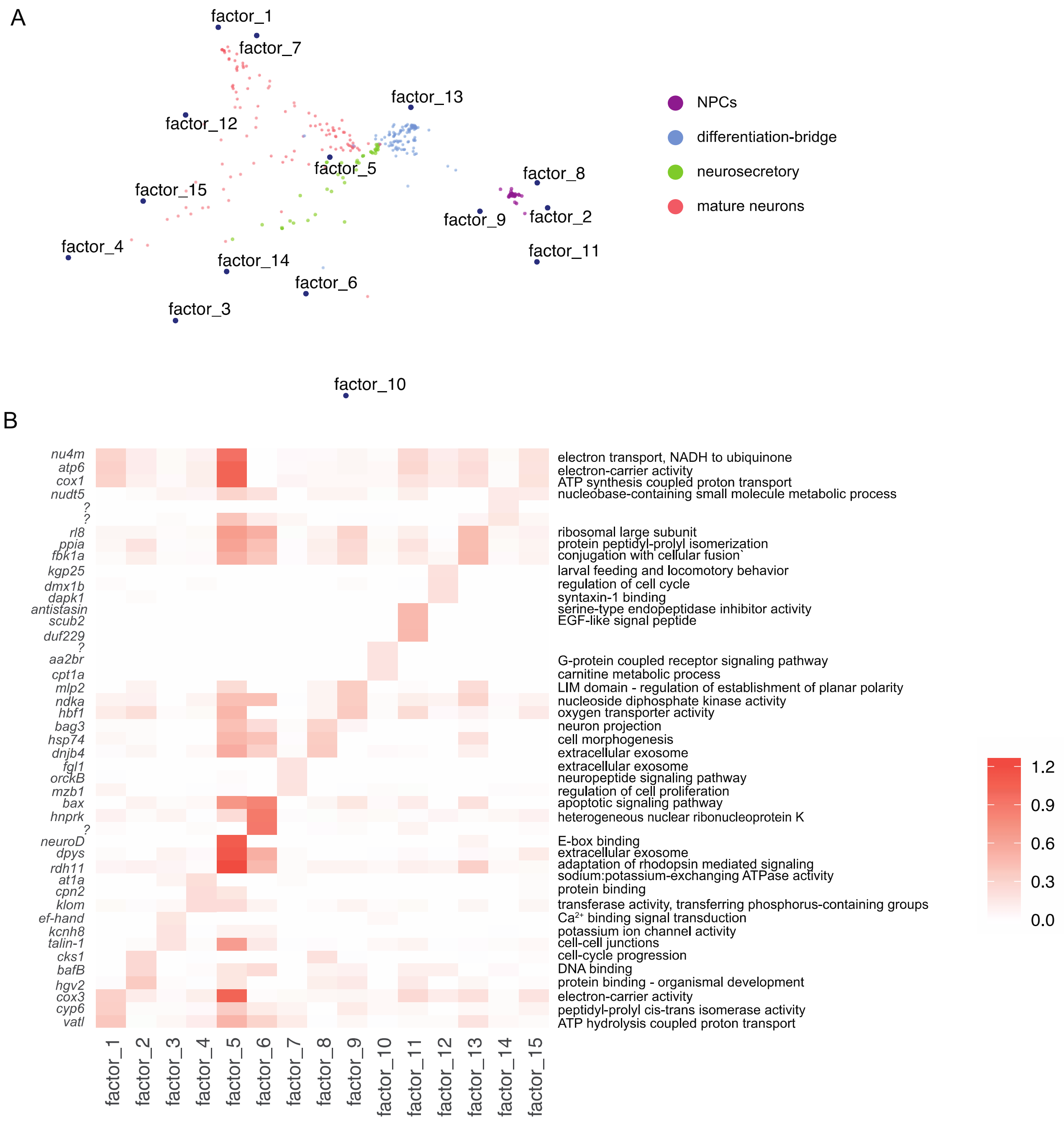


**Figure S8: SWNE visualizations of the stage 5 neural subcluster.** (A) SWNE (k = 20) representation of stage 5 neural cells with clusters labeled by cell type identities and gene modules represented as factors embedded within the SWNE visualization. (B) Heatmap showing the top factors ($p < 0.01$) identified with SWNE analysis using GO-term based gene sets that vary across all neural cells. The GO terms associated to each factor are shown on the right. Each row represents a gene and each column represents a factor expressed in a particular cell type based on the SWNE embeddings shown in A.

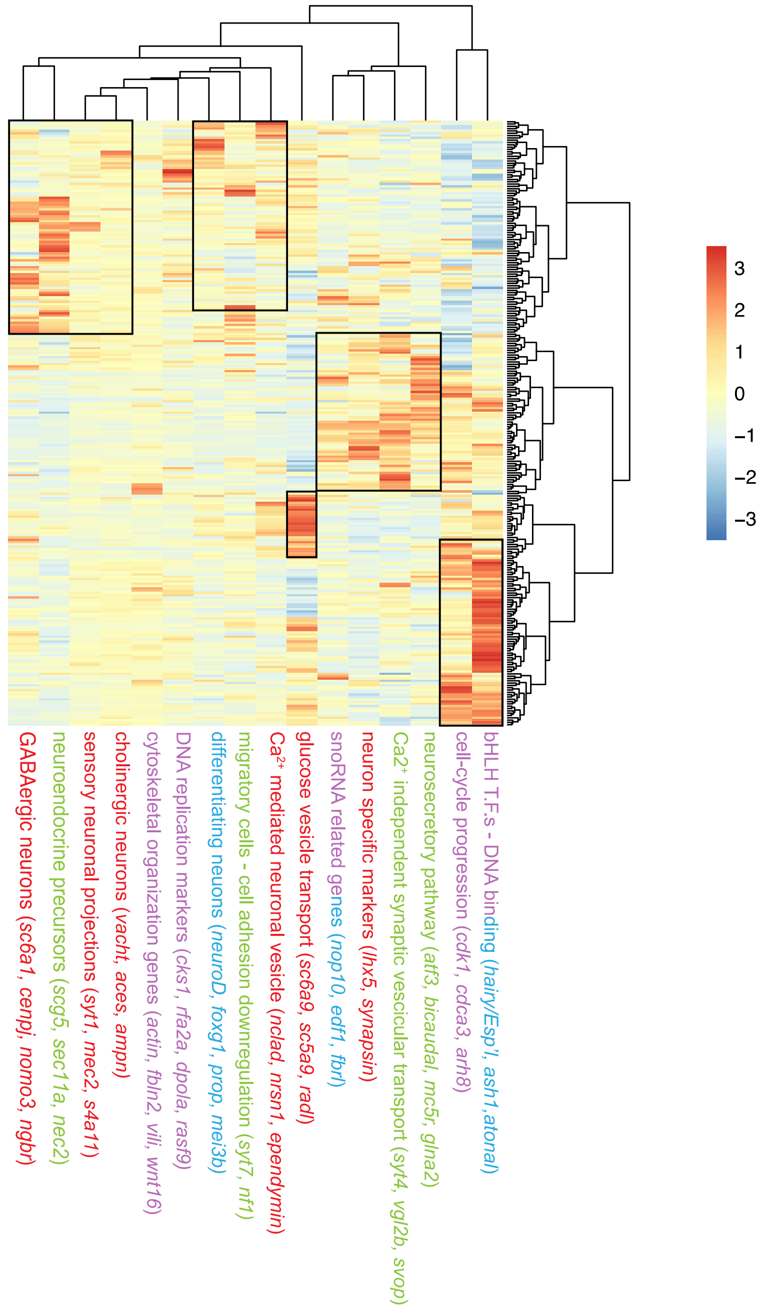


**Figure S9: Heatmap showing coregulatory gene modules within the stage 5 neural subcluster generated by Monocle3.** Individual rows indicate single cells whereas columns indicate clustered gene modules coregulated within each cell. Categories of markers and example marker genes are shown adjacent to the heatmap. See Fig. 7 for the neural differentiation trajectory generated using this heatmap. The expression bar on the right indicates upregulation and downregulation of genes in each module (blue: downregulated, red: upregulated). The hierarchical clustering output from Monocle recapitulates our observations from SWNE analysis (Fig. S8) and our Seurat clustering. The color of the individual GO terms corresponds to the color of the cell-cluster in Fig. 6A where these gene-modules were detected.

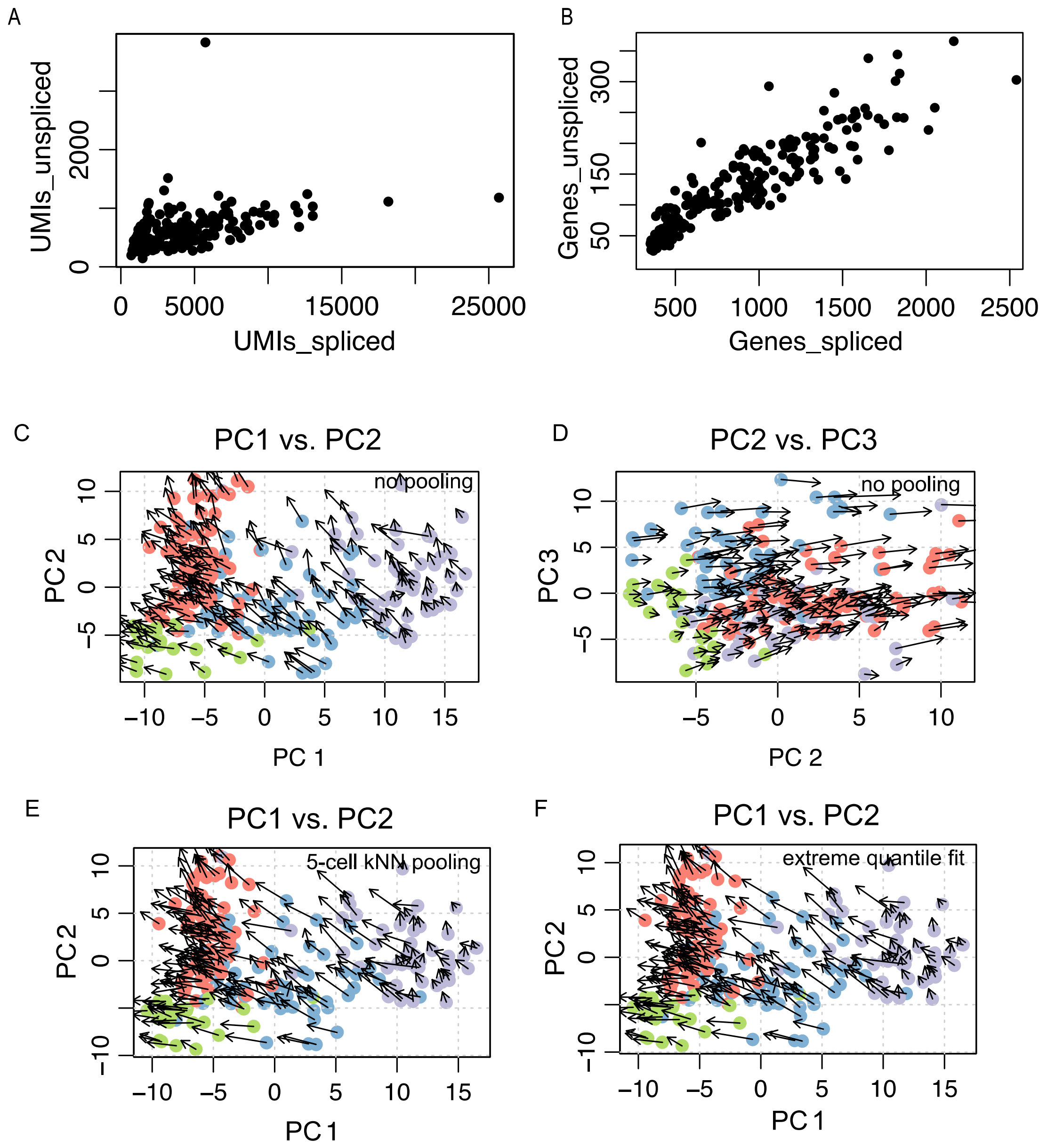


**Figure S10: PCA visualization of neural cell type RNA velocities.** (A) Scatterplot shows number of UMIs associated with spliced reads on x-axis versus UMIs associated with unspliced reads (B) Scatterplot shows number of genes associated with spliced reads with that associated with unspliced reads. (C, D) Projections on the first three principal components are shown. Please refer to Fig. 5, 6 and 7 for t-SNE, UMAP and RNA velocity visualizations of the cell types and the annotations of the subpopulation. The velocity was estimated using gene-relative fit for individual cells (i.e. without cell or gene pooling). Overall PC1 captures main neural cell type differentiation axis while PC2 captures separation between NPCs and other neural cell types. PC3 separates the neural differentiation bridge (blue) from the other cell types. (E) Gene-relative velocity estimates shown with k=5 cell kNN read pooling. (F) Velocity estimates with gamma slope and offset fit using only cells within the top or bottom 2% expression quantile for each gene. Cell kNN pooling of k=5 was used in calculating the estimates.

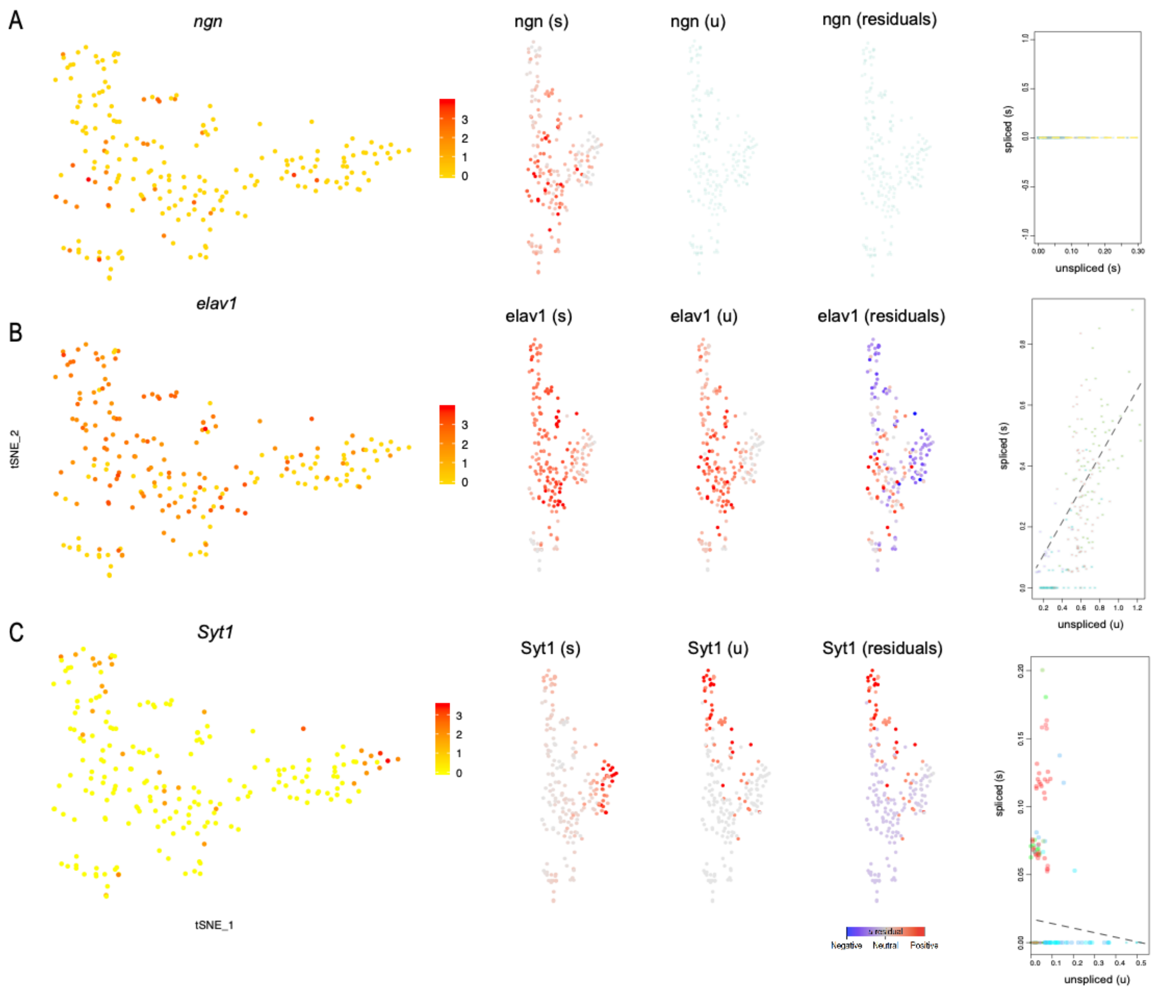


**Figure S11: Dynamics of RNA turnover during neural differentiation.** (A–C) A selection of phase-portrait plots that show genes underlying the observed velocity field as shown in Fig. 7. Markers for NPCs (A), differentiating bridge (B) and mature neurons (C) displayed above. Out of the three, no detectable velocity was observed for *Ct-ngn* (A). *Ct-elav1* (B) showed positive velocity while *Ct-syt1* (C) showed negative velocity. In each panel A, B and C, the first column highlights expression of individual genes in t-SNE space. The second column represents the observed expression profile of individual spliced molecules while the third column for each gene represent that for unspliced molecules. The fourth column illustrates the magnitude of the residuals (i.e. the difference between observed and expected unspliced abundance that tracks with the velocity estimation). The fifth column (boxes) show the phase-portraits and fits of the equilibrium slope (gamma-fit) for the neural cells at stage 5. For each gene, a spliced-unspliced phase-portrait is shown with the dashed line representing the gamma fit.

#### Supplementary Tables

**Table S1**: General statistics of sequencing and alignment as an output of the Cell Ranger pipeline. Additional metrics shown in the last four rows were derived from the Seurat R package.

| General | ST4pt | ST5 |
|---|---|---|
| Estimated number of total cells | 34,592 | 16,434 |
| Mean Reads per cell | 7,251 | 17,837 |
| Median genes per cell | 174 | 241 |
| **Sequencing** | | |
| Number of Reads | 250,821,573 | 293,135,282 |
| Valid Barcodes | 98.1% | 98.0% |
| Valid UMIs | 99.9% | 99.9% |
| Sequencing Saturation | 58.1% | 53.0% |
| Q30 bases in barcode | 94.8% | 94.9% |
| Q30 bases in RNA read | 83.7% | 84.7% |
| Q30 bases in Sample Index | 94.4% | 94.3% |
| Q30 bases in UMI | 93.6% | 93.7% |
| **Mapping** | | |
| Reads mapped to genome | 92.0% | 92.5% |
| Reads mapped confidently to genome | 67.1% | 67.5% |
| Reads mapped confidently to intergenic regions | 11.8% | 12.2% |
| Reads mapped confidently to intronic regions | 7.2% | 6.8% |
| Reads mapped confidently to exonic regions | 48.1% | 48.5% |
| Reads mapped confidently to transcriptome | 41.7% | 42.6% |
| Reads mapped antisense to gene | 0.1% | 0.1% |
| **Cells** | | |
| Estimated number of cells | 34,592 | 16,434 |
| Fraction Reads in cells | 59.3% | 42.0% |
| Mean reads per cell | 7,251 | 17,837 |
| Median genes per cell | 174 | 241 |
| Total genes detected | 16,668 | 17,186 |
| Median UMI counts per cell | 740 | 1,145 |
| **Sample** | | |
| Transcriptome | Capca1_genbank | Capca1_genbank |
| Chemistry | Single Cell 3’ v3 | Single Cell 3’ v3 |
| Cell Ranger version | 3.1.0 | 3.1.0 |
| **Seurat** | | |
| Seurat version | 3.1.4 | 3.1.4 |
| R version | 3.5.2 | 3.5.2 |
| Cells provided as input | 34,592 | 16,434 |
| Cells post QC | 1072 | 1785 |
